# The Life History Of Human Foraging: Cross-Cultural And Individual Variation

**DOI:** 10.1101/574483

**Authors:** Jeremy Koster, Richard Mcelreath, Kim Hill, Douglas Yu, Glenn Shepard, Nathalie Van Vliet, Michael Gurven, Hillard Kaplan, Benjamin Trumble, Rebecca Bliege Bird, Douglas Bird, Brian Codding, Lauren Coad, Luis Pacheco-Cobos, Bruce Winterhalder, Karen Lupo, Dave Schmitt, Paul Sillitoe, Margaret Franzen, Michael Alvard, Vivek Venkataraman, Thomas Kraft, Kirk Endicott, Stephen Beckerman, Stuart A. Marks, Thomas Headland, Margaretha Pangau-Adam, Anders Siren, Karen Kramer, Russell Greaves, Victoria Reyes-García, Maximilien Guèze, Romain Duda, Álvaro Fernández-Llamazares, Sandrine Gallois, Lucentezza Napitupulu, Roy Ellen, John Ziker, Martin R. Nielsen, Elspeth Ready, Christopher Healey, Cody Ross

## Abstract

Human adaptation depends upon the integration of slow life history, complex production skills, and extensive sociality. Refining and testing models of the evolution of human life history and cultural learning will benefit from increasingly accurate measurement of knowledge, skills, and rates of production with age. We pursue this goal by inferring individual hunters’ of hunting skill gain and loss from approximately 23,000 hunting records generated by more than 1,800 individuals at 40 locations. The model provides an improved picture of ages of peak productivity as well as variation within and among ages. The data reveal an average age of peak productivity between 30 and 35 years of age, though high skill is maintained throughout much of adulthood. In addition, there is substantial variation both among individuals and sites. Within study sites, variation among individuals depends more upon heterogeneity in rates of decline than in rates of increase. This analysis sharpens questions about the co-evolution of human life history and cultural adaptation. It also demonstrates new statistical algorithms and models that expand the potential inferences drawn from detailed quantitative data collected in the field.

## 1. INTRODUCTION

As a slow-developing primate, humans exhibit puzzling life history traits. Primates in general, and especially the apes, have slow life histories, with late age of first reproduction and singleton births. But even compared to other hominoids, humans have longer child-hoods, shorter inter-birth intervals, and extended post-reproductive lifespans (Jones 2011). That is, human children are slower to develop and more dependent, but we nonetheless have more of them, more quickly. These traits are plausibly unique to the genus *Homo*, but the timing and adaptive origins of the human life history strategy remains unsettled (Schwartz 2012).

One way for humans to ease the costs of expensive childhoods is through alloparental investments from highly productive adults (Kramer 2010). There are at least two major questions lurking within, however. The first is: Which individuals provide allocare? Any answer to this question will have implications for how selection operates on other aspects of life history. The second: Is childhood itself more than just a period required for growing large and physically adept? Is it also required for individuals to learn complex, culturally-evolved skills (Gurven et al. 2006)? What role does childhood play in the cultural evolution of complex, productive skills in the first place (Henrich and McElreath 2003)?

Any satisfactory model of human life history must address the integration of growth, reproduction, cognitive development, skill development, sociality, and cultural evolution. This is not easy. As a result, existing models make progress by omitting some features. The most advanced attempt we know is the optimal control model of González-Forero et al. (2017). While this model omits cultural dynamics for acquired skills, it does successfully integrate growth, cognitive and skill development, and reproduction in overlapping generations. By solving for the optimal life history, the model suggests natural selection for delayed growth, early investment in cognition, and delayed reproduction. The brain gets big first, and only then the body, because this allows a longer window of learning and ultimately higher adult productivity. These results are similar to the *embodied capital hypothesis* (Kaplan et al. 2000), in which highly productive foraging and food sharing by adult men supports alloparental investments in offspring. From this point of view, human life history traits stem from the highly complex human foraging niche, which selects for delayed maturation by requiring an extended period of learning before adults are able to achieve high productivity. In contrast, Hawkes et al. (1998) emphasize provisioning of grandchildren by post-reproductive women, which selects for longer lifespans. This perspective sees childhood as a consequence of prolonged lifespan, not a trait that needs to be explained as having its own direct function (Charnov 1993). A spectrum of models exists, in which adult foraging is variably influenced by size, skill, and culturally-transmitted knowledge, and different amounts of time are needed for individuals to acquire and perfect adult skills.

To develop and test models, anthropologists have used observational studies of subsistence hunting, with a focus on variation across the lifespan. For example, Walker et al. (2002) and Gurven et al. (2006) report data from the southern Neotropics that subsistence hunters achieve high proficiency only after reaching advanced ages, roughly 35 to 45 years old. Because hunters achieve adult size and strength much earlier in life, these results are consistent with the embodied capital hypothesis and its emphasis on the gradual mastery of cognitively complex hunting strategies. But comparative data from other contexts have been scarce. Among the few other empirical studies, some find slow skill development (e.g., Ohtsuka 1989) while others do not (Bird and Bliege Bird 2005).

More and better estimates of age-related foraging skill are necessary inputs into all evolutionary models of human life history. Associations between brain development, cultural knowledge, physical skill, and foraging performance at each age constrain the models we specify: quantitative and representative estimates of these variables are needed to parameterize optimal life history models like González-Forero et al. (2017). Variation across individuals informs models of food sharing and other investments, both within and between generations. Variation across sites and contexts informs models of tradeoffs and how individuals cope with them.

In principle, skill and production in other subsistence economies is equally relevant to understanding human life history. Garden production and animal husbandry depend upon the same cognitive and developmental foundations as hunting and gathering. We focus on subsistence hunting for two reasons. First, the data are easier to model than are gardening and herding—hunting returns are easier to identify with specific individuals and labor allocations. Second, hunting is practiced, to some extent, everywhere. It is both a primitive economy and a modern one that has endured the emergence of other subsistence strategies. The breadth of hunting in diverse ecological settings provides a compelling range of evidence.

Studies of hunting returns are nevertheless inferentially challenging. A typical outcome variable, such as kilograms of harvested meat, may be a mixture of zeros and skewed positive values that violate assumptions of conventional regression models (McElreath and Koster 2014). The available foraging data often exhibit imbalanced sampling of individuals and age groups. Predictor variables may be missing or measured with uncertainty. These problems are surmountable in any individual study, but comparative inferences are challenging when studies rely on heterogeneous statistical solutions.

In this paper, we address the inferential and comparative challenges within a novel statistical framework. We assemble the largest yet data base of individual human hunting records, comprising over 21,000 trips from 40 different study sites. These data elucidate the extent to which the ontogeny and decline of hunting skill are attributable to individual level or site-level factors, and the comparative analysis help to mitigate over-generalization from individual studies. The results of this study consequently inform subsequent theorizing about the evolution of life history traits in humans.

Our statistical approach accepts the imperfections of the sample and conservatively pools information, both among individuals within sites and among sites within the total sample. The goal is not to substantiate any particular theoretical model of human evolution, nor to pretend that the data are sufficient for all inferential objectives. Rather, the goal is to show what can be inferred froma statistical approach that uses all available data and treats missing data and measurement error conservatively. One of the most important aims is to highlight the limits of existing data and approaches so that future empirical and inferential projects can make further progress.

Our analysis supports the general conclusion that skill peaks between 30 and 35 years of age, well after the age of reproductive maturity. Peak skill is typically not much higher than skill during early adulthood, however. Declines with age are typically slow—an average 56 year old has the same proportion of maximum skill as an average 18 year old. There is considerable variation both among sites and individual hunters within study sites. Variation among individuals is described more by heterogeneity in the rate of decline than the rate of gain. Partly owing to heterogeneous data collection methods across sites and anticipated biases from omitted variables, not much can be inferred yet from this sample and model about the causes of differences among sites.

These results are computed conservatively, but as with any analysis, the results are dependent on the model and sample. In the future, alternative statistical approaches may enhance the inferences provided by the present study. Consequently, we are careful to describe both the nature of the model and the data, and all of the code and data are publicly available.^1^ Our objective is to involve more theorists and empiricists in the long-term project of constraining and informing models of how life history is integrated with human behavioral adaptation.

## 2. DESCRIPTION OF THE DATA

The total sample contains 1,821 individual hunter, 23,747 hunter-level outcomes, and 21,160 trips across 40 study sites (Figure 1). To compile the dataset, the first author searched for relevant studies on subsistence hunting in the anthropological and biological literature, subsequently contacting authors to invite them to contribute data. The contributors submitted data in a standardized format that included variables for the biomass acquired on terrestrial hunting trips, the ages of the hunters at the time of the hunt, the duration of the trip, the hunting weaponry carried by the hunters, and the presence of dogs or assistants (e.g., porters). Our data are restricted to hunting, and exclude gathering, because of the paucity of data on gathered plant foods.

**FIGURE 1.**
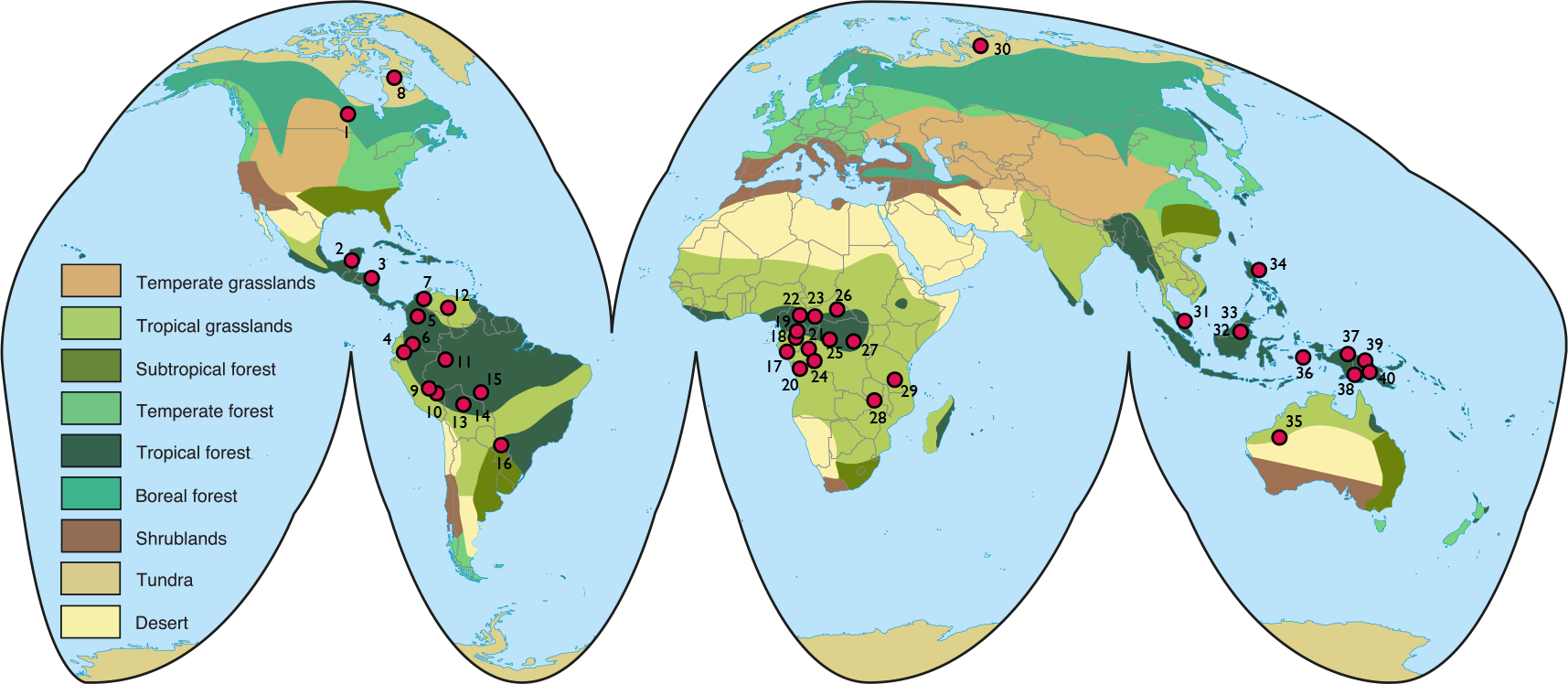
Distribution of study sites. For the key, see Table 1.

There is tremendous imbalance in sample size across units. One site contributes only 6 trips from 2 individuals. Another contributes more than 14,000 trips from 147 individuals. Some individuals contribute only a single outcome, while others contribute dozens. The majority of the sample comprises male hunters, with too little data on female hunters to infer generalizable sex differences. (This does not imply that men’s production and skill is more relevant to human evolution, nor that women’s foraging skill would necessarily exhibit either the same or a different functional relationship with age.) Most sites contribute primarily cross-sectional data, while a few others exhibit impressive time series. The statistical framework is designed to make use of all these data.

**TABLE I.**
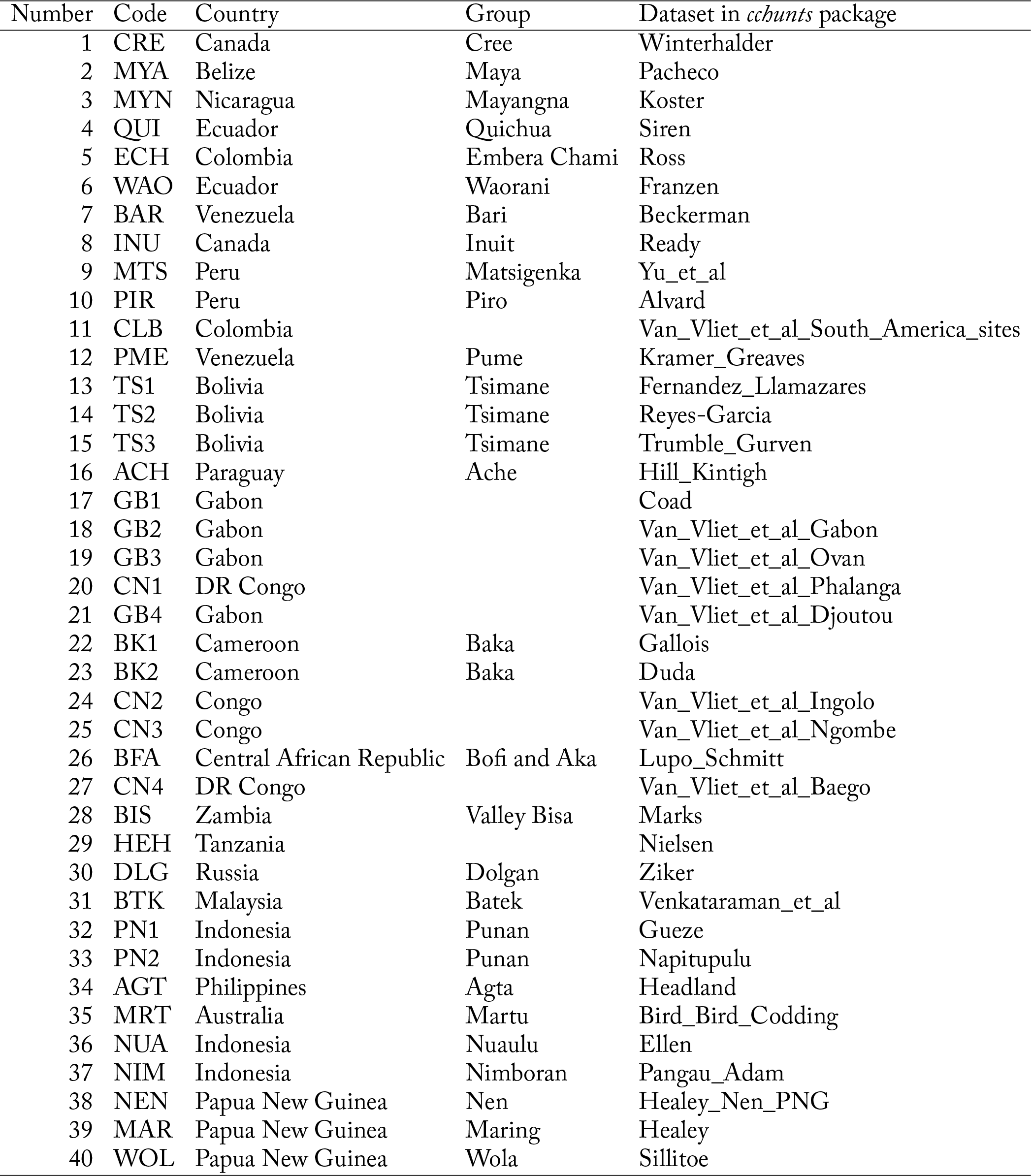
Study sites and their numerical and text codes. See the help file of the *cchunts* package for related citations.

## 3. THE LIFE HISTORY FORAGING MODEL

Since skill cannot be directly observed, what is required is a model with latent age-varying skill. This unobservable skill feeds into a production function for observable hunting returns. In this section, we define a framework that satisfies this requirement. We explain it one piece at a time, with a focus on the scientific justification. The presentation in the supplemental contains more mathematical detail, and the model code itself is available to resolve any remaining ambiguities about the approach. Our framework was developed and reviewed in the initial grant proposal (NSF #1534548) prior to seeing the assembled sample. Therefore, whatever the model’s flaws, they do not include being designed specially for these observations or chosen to produce a desired result.

One advantage of the latent skill approach is that it allows us to use different observations from different contexts—both solo and group hunting, for example—to infer a common underlying dimension of skill. But modeling even the simplest foraging data benefits from this approach, as hunting returns often are highly zero-augmented. Separate production functions for zeros and non-zeros are needed to describe such data. In principle, more than one dimension of latent skill could be modeled. We restrict ourselves to only one in the current analysis. With more detailed data, describing additional dimensions should be possible.

We implemented the model both as a forward simulation and as a statistical model. The forward simulation generates data with known parameter values, which are used to confirm that the estimated statistical model can recover the parameters. The code is available as part ofthe `cchunts` R package.

### 3.1. Latent skill model

One of the simplest life history models is the von Bertalanffy (1934) asymptotic growth model. We use this model to represent the increasing components of hunting skill as a function of age. These increasing components include knowledge, strength, cognitive function, and many other aspects that contribute to hunting success and increase but decelerate with age. For convenience, label the composite of these components *knowledge*. Assume that the rate of change in knowledge with respect to age *x* is given by d*K*/d*x* = *k*(1 − *K*(*x*)). This means only that knowledge increases at a rate proportional to the remaining distance to the maximum—the more there is left to learn, the more one learns. Solving this differential equation yields the age-specific knowledge of a hunter at age *x*:

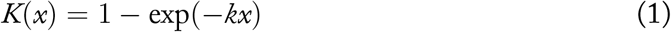

where *k* > 0 is a parameter that determines the rate of increase. To account for senescence, we assume that production capacity *M* declines at a constant rate, given by d*M*/d*x* = −*mM*(*x*). Solving this yields:

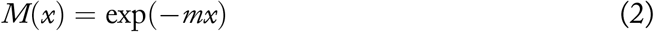

where *m* > 0 represents the rate of decline. The total age-specific skill is given by a weighted product of these two functions:

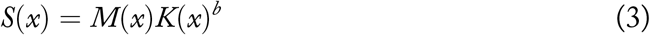

where the parameter *b* controls the relative importance of *K*. In economic terms, *b* is the knowledge elasticity of skill. We assume that *k* and *m* may vary across individuals—some people learn faster or senesce more slowly—while *b* is a property of the production context at a given study site.

This model is among the simplest we can construct. Nevertheless, it is capable of describing diverse age-specific skill curves. Figure 2 illustrates the general shapes of each component of the model, as well how variation in each component may produce variable life histories. Each plot in this figure shows age on the horizontal axes. The top row of the figure illustrates the general shape of each component (left and middle) and one possible resulting lifetime skill curve (right). The bottom row shows 10 different, random knowledge and senescence curves, with their implied random skill curves. These demonstrate that even a model as simple as this one, with only three parameters, is nevertheless capable of producing many diverse age-specific curves. This approach brings two more advantages, as compared to the use of polynomial functions of age. First, the parameters have straightforward biological interpretations. Second, these functions do not exhibit instabilities such as Runge’s phenomenon (Runge 1901) that complicate fitting and prediction.

**FIGURE 2.**
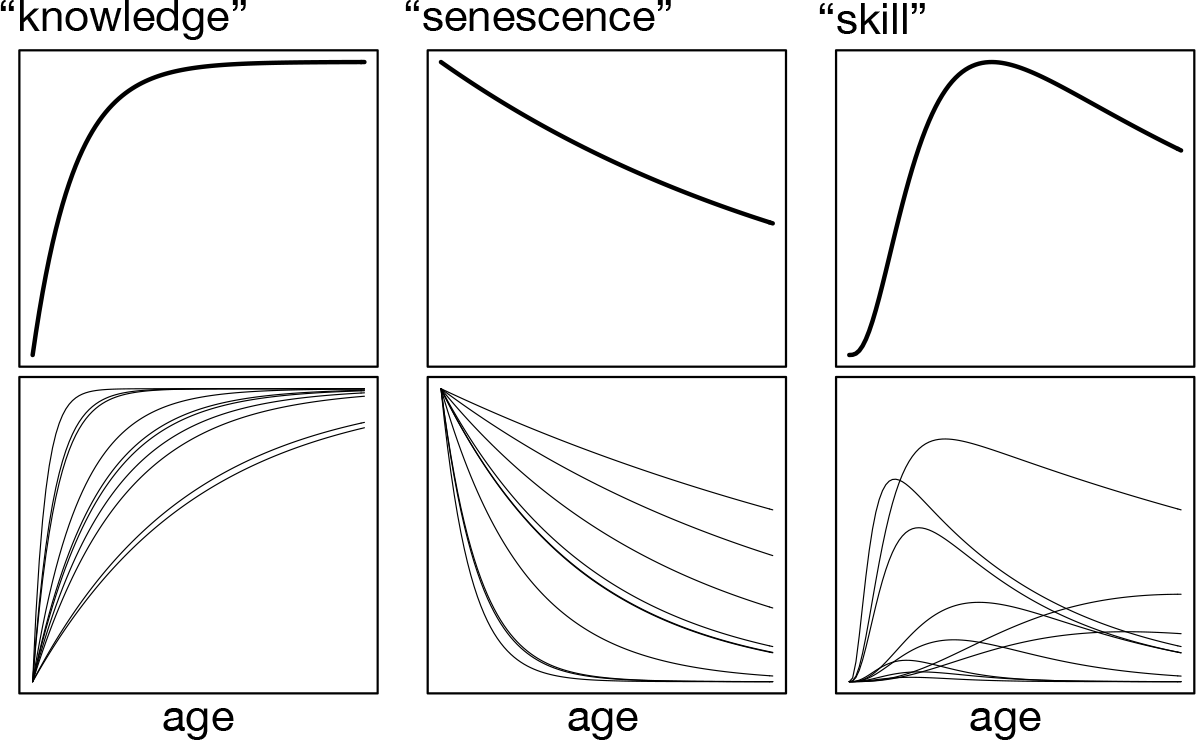
The age-specific skill model. Top row: Increasing components, “knowledge,” anddecreasingcomponents, “senescence,” multiplytoproduce relative productive potential at each age, “skill.” See the supplemental material for equations. Bottom row: Variation in the components combines to produce a diverse array of possible skill functions.

These functions also have clear weaknesses. Neither the rate of gain *k* nor the rate of loss *m* is plausibly constant over large age ranges. The rate of variation in body growth, for example, will produce rate variability in skill growth. And near the end of life, skill loss should accelerate rather than slow down. Although the data analyzed in this paper do not span the age ranges in which this variation would occur, we should be cautious about overgeneralizing from this analysis.

The final component of the core skill model is partial pooling of information. Since these data contain repeat measurements on the same units—individuals and sites—as well as substantial imbalance in sampling of these units, partial pooling via multilevel modeling provides superior estimates. The variation among individuals is also a target of inference. We employ two levels of hierarchical pooling (Figure 3). First, the life history parameters *k* and *m* are pooled across individuals within each site (left column, Figure 3). In standard terminology, *k*_ID_ and *m*_ID_ for each individual are random effects drawn from a bivariate distribution. Each site also has its own value for *b*, reflecting variation in the relative importance of knowledge across sites. Therefore each site has its own distribution of skill functions (middle column). Finally, the site distributions are pooled together to regularize inference at the second level (right column), producing a distribution of site distributions.

**FIGURE 3.**
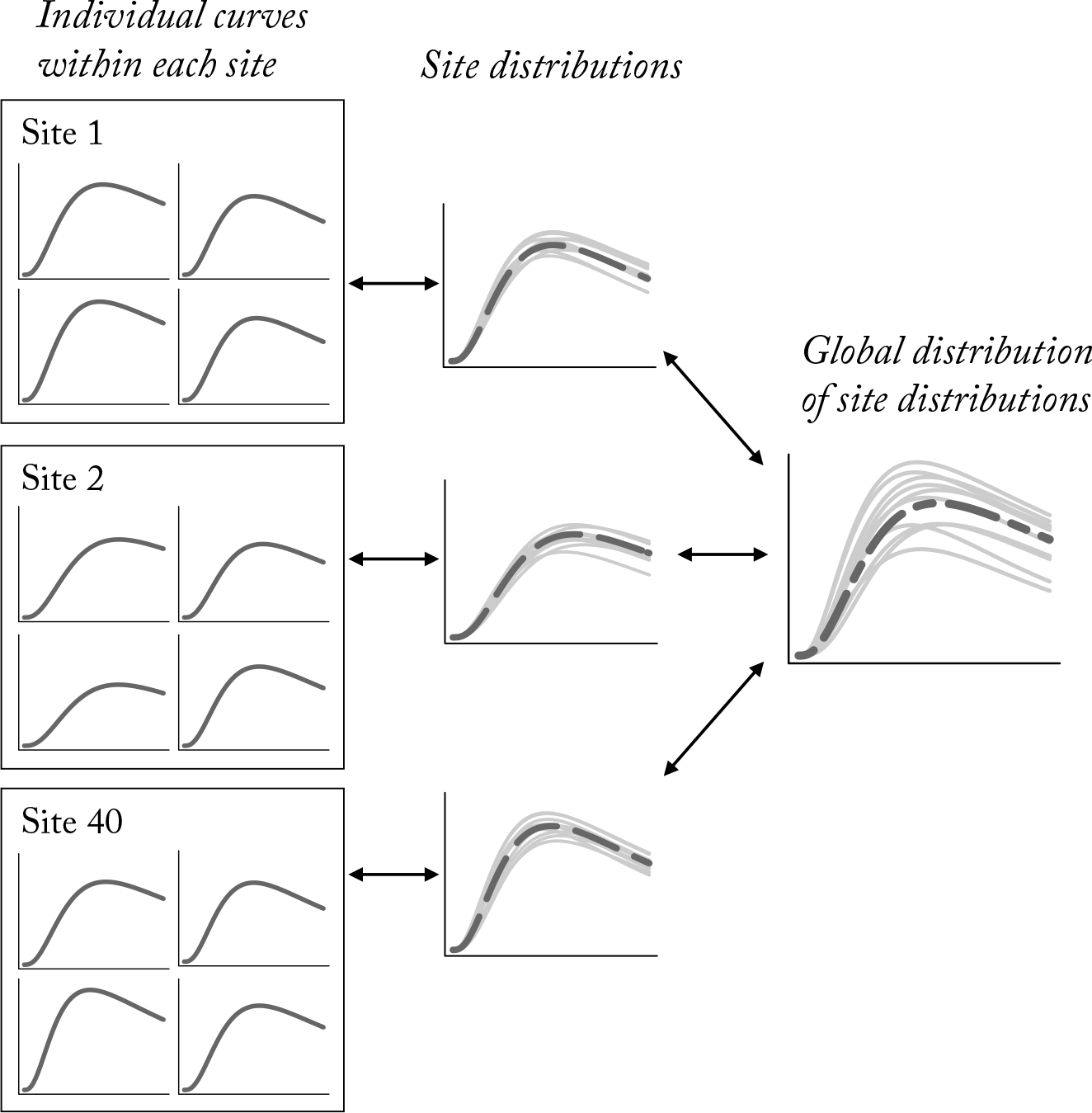
Hierarchical structure of skill functions within the inferential model. Within each site (left) a skill curve is inferred for each hunter. Individuals within each site are pooled using a distribution of individual skill curves (middle). Finally, the distributions of parameters within each site are again pooled using a distribution of distributions (right). This formulation allows variation among individuals to vary by site.

To an extent, this global distribution is a statistical fiction that is necessary to pool information properly among sites. However, it is also a target of inference, providing a weighted summary of all of the evidence across sites.

### 3.2. Production model

Skill is not directly observable. Rather, we must infer it by its effects on hunting productivity. This requires introducing a layer of production functions through which skill acts. The production data available to us contain two correlated components: (1) the probability of a successful trip that produces a non-zero harvest and (2) the size of harvests obtained on successful trips. We model each with a standard log-linear function of labor, skill, and technology. Specifically, for successful trips, the mean expected harvest at skill *S* is given by:

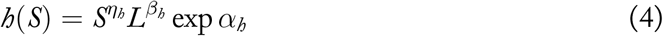

where *η*_*h*_ is the elasticity of skill, which determines the magnitude of skill differences on harvest, 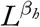 is the labor allocated with its elasticity *β*_*h*_, and *α*_*h*_ is a linear model including terms for technology and cooperation variables. Notice that harvest increases with both skill and labor, but that the elasticity of each determines the impact of any increase. The full distribution of harvests is assumed to follow a gamma distribution, which allows for the highly skewed distributions typical of many hunting data sets. We have used this assumption in previous work (McElreath and Koster 2014). However, a log-normal distribution of harvests would work as well. The important features are to impose a zero lower bound and to allow for positive skew. If we had detailed data on the encounters and pursuits of individual prey types, we could build a mixture distribution to better describe observed harvest sizes. But such data are available in very few cases. For comparability across sites and compatibility with the logit function described next (equation 5), we have proportionally standardized harvests relative to the maximum harvest size at the respective study sites.

A similar approach provides a Bernoulli distribution of success/failure. The probability a trip produces a non-zero harvest is:

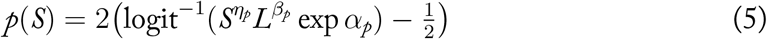

The terms enclosed within the interior parentheses recapitulate the log-linear production function of the above equation (4). The remainder of the function re-scales the log-linear model so that *p*(*S*) varies continuously from zero to one and *p*(0) = 0.

This is a descriptive approach. It has the advantage of being able to describe many possible relationships between skill, labor, and technology. Figure 4 illustrates some of the model’s features. Each plot in this figure shows labor input—hours allocated to foraging—on the horizontal axis. From left to right, the plots show the probability of a non-zero harvest, the expected harvest size on a successful trip, and the expected returns resulting from the product of the two. Each row illustrates the impact of one type of variation—variation in individual skill in the top row and variation in hunting group size in the bottom row. The first thing to notice is that the function implies monotonic returns to labor. Marginal returns must always either increase or decrease with labor. Second, skill and labor can influence hunting success and harvest size quite differently. There is no assumption that skill or labor is equally important for both components of production. And since technology can influence elasticity of skill and labor, technology can have independent effects as well.

**FIGURE 4.**
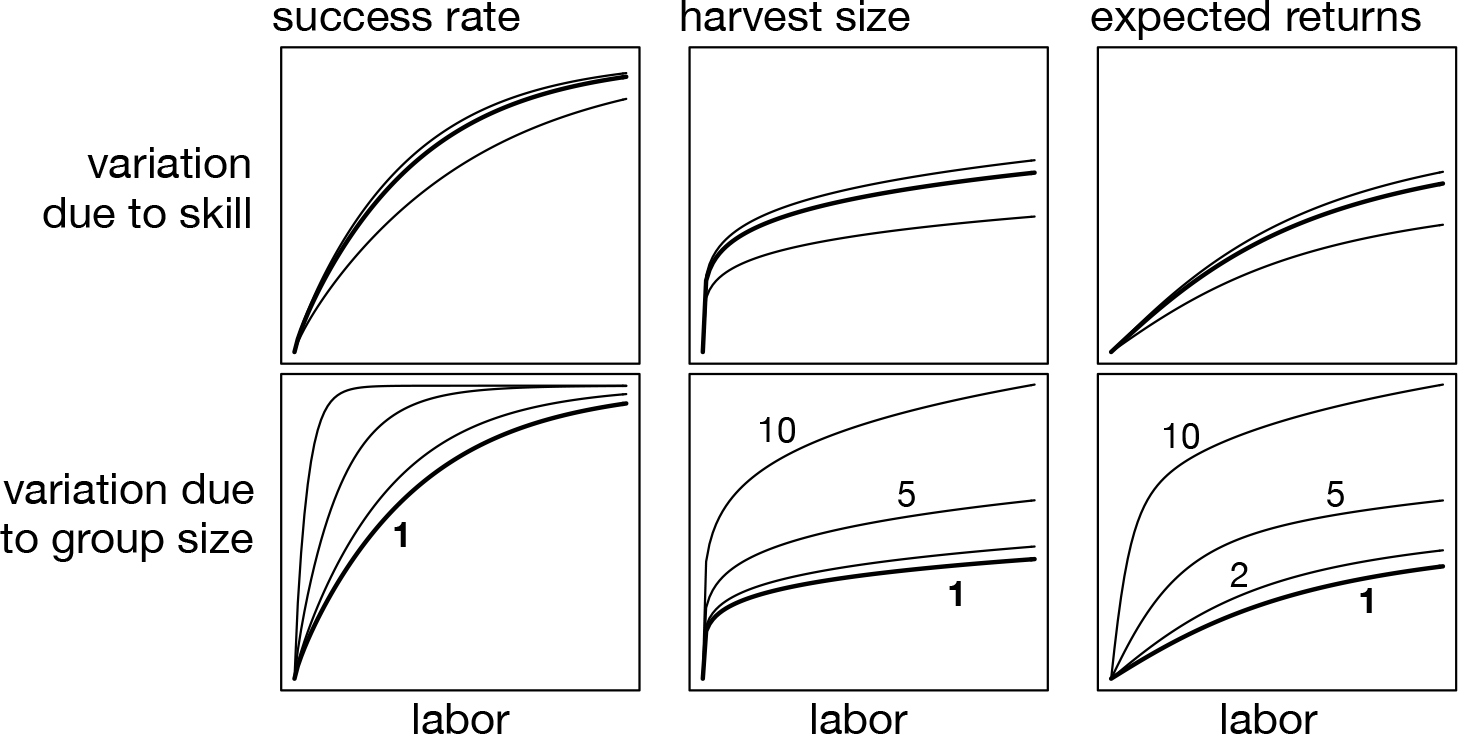
Example production functions for observed harvests. Expected harvest (righthand column) is the product of the probability of a non-zero harvest (lefthand column) and the expected size of a non-zero harvest (middle column). The top row shows how each component may vary with skill. The bottom row shows how each may vary with number of hunters.

### 3.3. Cooperative trips and aggregated harvests

Many of the hunting trips in our sample are cooperative, in the sense that multiple hunters of varying skill interact in producing returns. The harvests on these trips may be assignable to individual hunters or alternatively credited to the group as a whole. We handle cooperative trips by treating them as analogous to technology, with group size represented as a coefficient in the production equation. When returns are aggregated to the level of the group rather than assigned to individual hunter, we replace individual hunter skill in the production equation with average skill of the group.

### 3.4. Missing values and measurement error

Our sample embodies common statistical challenges. First, there are many missing values, notably for trip duration and the presence of dogs on trips. Second, there is measurement error, notably for individual ages. The customary solution to these problems is to drop all cases with any missing values and to replace uncertain measurements with their means. Instead of dropping cases with missing values, however, we model the unknown values. This allows Bayesian imputation of missing values, averaging over uncertainty in unobserved durations. We rely upon the same principle to handle measurement error in age. The co-authors who contributed datasets to our sample assigned a standard error to each recorded age. Within the model, each hunter’s date-of-birth is replaced with an unknown parameter with a prior centered on the recorded age and with standard deviation equal to the recorded standard error. In a few cases, no age is recorded for an individual. In those cases, we assign a vague prior that covers the entire range of observed ages. For more details on these techniques, see Chapter 14 of McElreath (2015).

### 3.5. Inference

The full model contains just under 28,000 parameters. A wry quotation comes to mind, attributed to John von Neumann, “With four parameters I can fit an elephant, and with five I can make him wiggle his trunk” (Dyson 2004). With 28,000 parameters, maybe we can animate an entire stampeding herd?

In principle, perhaps, but in this application, we cannot. The reason is that these parameters are not free parameters. Many of these correspond to missing durations and age uncertainties, and so contribute little fit to the sample. Many of the remaining parameters arise from the hierarchical structure of the life history model. These parameters do not make it easier to fit the sample, but rather harder. They reduce overfitting, by pooling information among sampling units. For the remaining parameters, we adopt regularizing priors that are more conservative than the implied flat priors of typical non-Bayesian procedures (see Chapter 6 of McElreath 2015, for aa explanation). We present a complete description of the priors in the supplemental. Having fit alternative parameterizations of the model, we believe the results that we present in the next sections are qualitatively robust to changes in priors and even the hierarchical structure of the model. To facilitate alternative estimates of model parameters, though, we provide our annotated statistical code in the supplement.

Inference for a model with so many dimensions remains a challenge. Optimization approaches can fail in high dimensions. In high dimensions, a typical draw can be very far from the mode, a phenomenon known as *concentration of measure*. The most common Markov chain Monte Carlo algorithms, such as Gibbs sampling, also fail in high dimensions, since they tend to get stuck in local neighborhoods and poorly explore the posterior distribution. A family of algorithms known as Hamiltonian Monte Carlo perform much better in these settings (Neal 2010, Betancourt 2017). We used Stan’s implementation of Hamiltonian Monte Carlo (Stan Development Team 2016) to sample from the posterior distribution of the full model. We present the results in the next section as summaries of 500 draws each from 10 chains. We assessed chain convergence and mixing efficiency by means of the Gelman-Rubin diagnostic 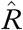 and an estimate of the autocorrelation adjusted number of samples, `n_eff`, both as calculated by`rstan` version2.16.1. We also visually inspected trace plots of the chains to ensure that they converged to the same target distribution. Finally, we compared the posterior predictions to the raw data to ensure that the model corresponds to descriptive summaries of the sample. All parameters demonstrate good mixing, with 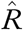 values below 1.01. Since there are more than 20,000 parameters, we cannot conveniently provide a table of these diagnostics. However, an R data file with samples and functions for diagnostics are available as an online supplement.

## 4. RESULTS

The are many ways to summarize the model inferences. We focus on three foundational issues that motivated the project.

1. What is the overall pattern of skill development?
2. How variable is this pattern within and between societies?
3. Which components of the model—increases early in life or declines later in life— describe variation?

We consider each of these issues in order.

### 4.1. Overall pattern

At the highest level of pooling, the model provides a statistical answer to the question, “What is a typical human life history of hunting skill?” This is very much an abstraction, one that attempts to factor away all the variation in production functions and associated elasticities to reveal an underlying, dimensionless skill function. It cannot say much about absolute levels of production, either within or between societies. But it can inform comparisons of relative skill at different stages of life.

The statistically average hunter in this sample peaks at 33 years of age (top-left plot, Figure 5). However, this peak is not sharp. At age 18, this fictional average hunter has 89% of maximum skill. And skill declines slowly, such that skill falls below 89% of maximum only after age 56. The blue shading around the posterior mean in this plot shows the entire posterior distribution, fading out to transparent as probability declines. The narrowness of this interval reflects high confidence about the global mean of this sample.

**FIGURE 5.**
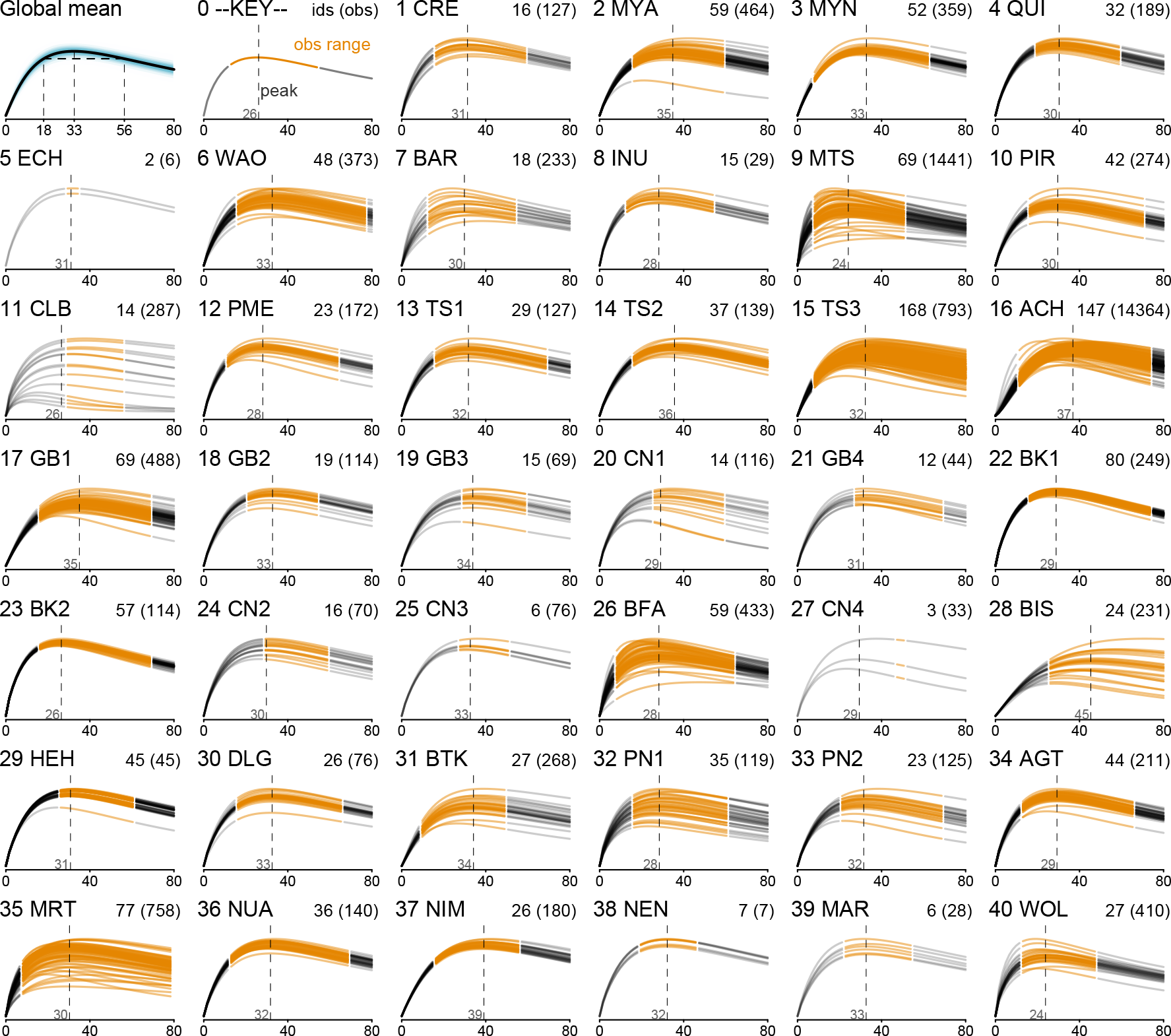
Global mean skill (top-left plot) and skill at each site. Each curve is the posterior mean skill for an individual. In the header of each individual plot, the site number and three letter code are shown along with the number of individual hunters in each sample, followed by the number of observed harvests in parentheses. The orange span of ages correspond to ages observed within each site, while the gray ranges were unobserved and are instead implied by the underlying model. The vertical dashed lines show the average ages at peak.

While the overall pattern is clear, not every site nor individual hunter exhibits the same pattern. The individual site plots in Figure 5 illustrate this variation. Each site displays the mean skill function for each hunter in the sample from that site, but note that the plots do not depict the uncertainty around individual skill curves. Nevertheless, there is strong evidence of high individual variation at some sites, such as the Matsigenka (9 MTS), the Aché (16 ACH), and the Martu (35 MRT). Differences among individuals can be quite large. Some individuals have half the adult skill of others in the same community.

At some sites, such as the Dolgan (30 DLG), the individual curves tend to cluster together around a central mean. This is not an indication that the model believes these hunters are all the same. Rather, there is not sufficient evidence to indicate that they are different, often stemming from relatively small sample sizes. In the supplemental, we present another version of this grid that instead shows simulated hunters sampled from the posterior distribution, which better represents the individual-level variation implied by the model (Supplemental Figure 7).

For each site, the figure also displays the age of peak skill for a statistically average hunter, as indicated by the vertical dashed lines. While these peak ages cluster around 30 years of age, there is some noteworthy variation. On the low end, the Matsigenka (9 MTS) and Wola (40 WOL) peak early, near 24 years of age. Note that the best hunters at these sites tend to peak even earlier, a trend that is also evident among the Barí (7 BAR). On the higher end, the Aché (16 ACH) and Valley Bisa (28 BIS) peak at 37 and 45, respectively, but with relatively slow declines.

These skill functions are inputs into site-specific production functions. This means that the relationship between age and skill is further modified by labor allocation and site-specific details like technology. In the supplemental material, we produce versions of the grid in Figure 5 that display the other components of production, such as failure rates. One feature of the production components is that variation can arise from different sources. In some settings, hunters are distinguished primarily by the frequency of unsuccessful hunts. In other settings, there is greater individual-level variation in the amounts acquired on successful hunts.

### 4.2. Structure of variation

Skill functions vary both within sites and between sites. Which components of skill contribute to this variation? To address this question, we examine the model parameters that measure variation in the components *k* (rate of increase) and *m* (rate of decline) of the skill function. Since this is a non-linear model, we cannot exactly partition total variance. The impact of variation in a component of skill depends upon the values of all the other components (Goldstein et al. 2002). We can, however, consider relative sizes of components of variation on the latent scale.

First, we find greater variation in *m* than *k* within sites (top-left plot, Figure 6, cyan density). Between sites (orange), both *m* and *k* contribute about equally. This implies that while variation among hunters within each site is explained more by heterogeneous senescence, both components are important to variation across sites. Some caution is necessary here, since the relationship between *m* and *k* is not additive. However, the implication is that skill functions vary more late in life than early in life. This comports with the posterior means visualized in Figure 2.

**FIGURE 6.**
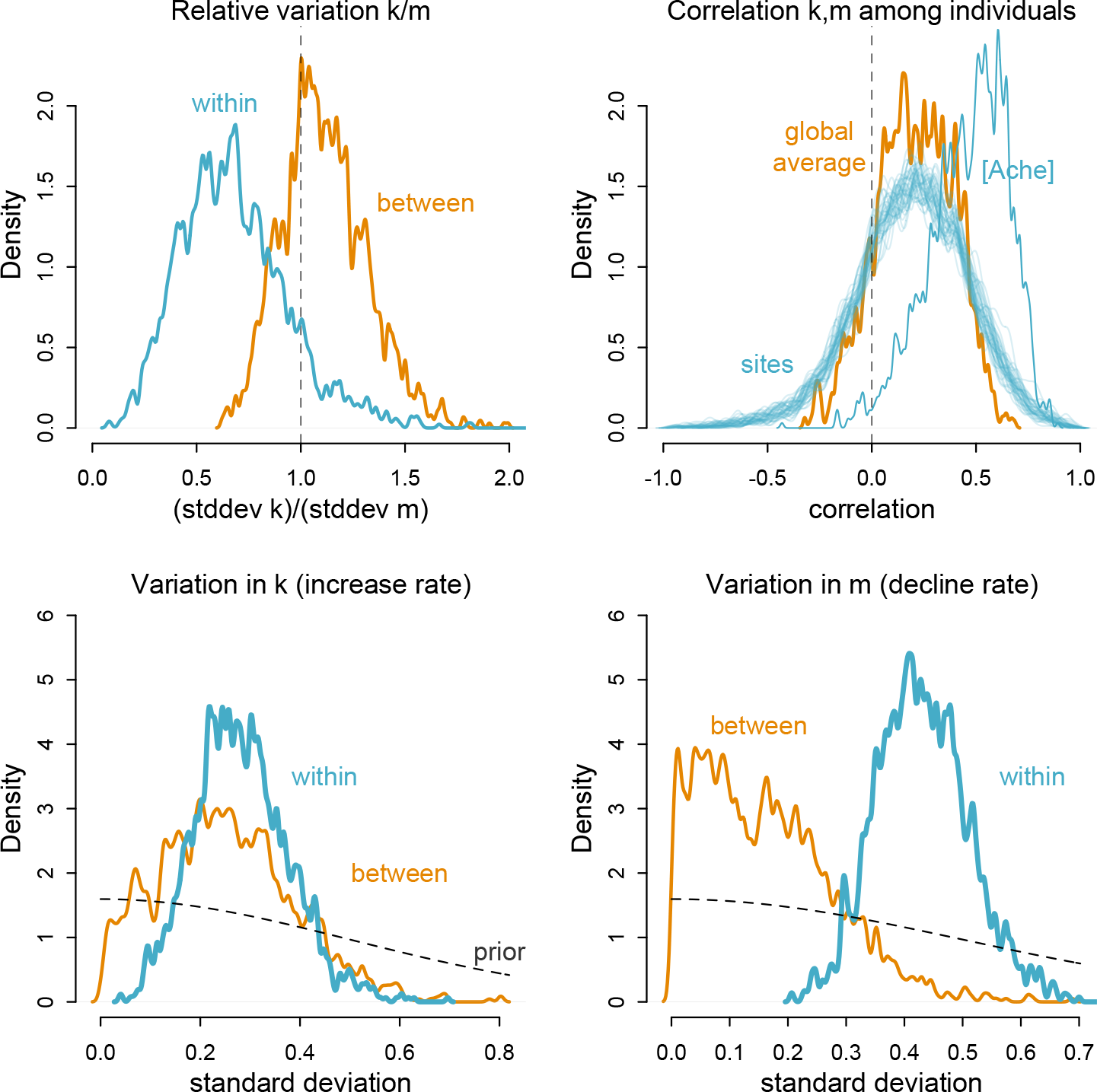
Variation in components of skill. Top-left: Relative variation in *k* and *m*. Horizontal axis is the ratio of the standard deviation of *k* to the standard deviation of *m*. The vertical line at 1 indicates equality of variances. Orange density is between site variation. Blue density is within site variation. There is more variation in *m* than *k*, both between and within sites, but the difference is much greater with insites, where *m* contributes more to variation among individuals than does *k*. Top-right: Correlation between *k* and *m* among individuals within sites. The orange density is the global average. Each blue density is a single site. The Aché stand out and are labeled separately. Bottom-left and bottom-right: Variation in *k* (left) and *m* (right) comparing within and between sites.

We also find a modest positive correlation between *k* and *m* (upper-right plot), suggesting that hunters who develop skill relatively quickly also show reduced declines in skill at higher ages. Each density is the upper-right plot is the posterior correlation between *k* and *m* for an individual site. This correlation is particularly pronounced for the Aché and modest otherwise. This may reflect the lack of longitudinal data on individual hunters at most study sites, limiting what can be learned about this correlation. In contrast, the Aché site contains enough time series data on individuals to make stronger inferences about the correlation.

Finally, variation in *m* and *k* can be decomposed additively within and among sites. We show the posterior distributions of the standard deviations in both *k* (bottom-left) and *m* (bottom-right) in Figure 6. The cyan densities are the standard deviations within sites, which correspond to the plausible values for variation among individuals. The orange densities are the standard deviations between sites, corresponding to the plausible values for variation among site means. The dashed curves in both plots show the prior distributions, which were the same for both within and between components. Because the orange curves remain flatter than the cyan curves, these plots show that for both *k* and *m*, there is relatively less information about variation among sites than within sites. While there is a hint that variation between sites contributes more to variation in *k* while variation within sites contributes more to variation in *m*, no strong inferences can be drawn until more information is available for inferring the between-site variance.

## 5. Discussion

Hunting is not an easy task. It is demanding of physical stamina, bio-geographical knowledge, experience reading environmental cues, close observation of animal behavior, technical finesse in the construction and maintenance of equipment, and often the ability to collaborate with partners. Even the most skilled hunters in our sample often return home with nothing. A hunter might spend hours tracking a trail that goes cold. A hunter might pass lots of small prey but never encounter something large enough to be worth pursuing. A hunter could strike prey but subsequently fail to catch or locate it. A hunter might even run out of arrows or ammunition and be forced to return with nothing. Every failure case implicates physical skills, cognitive skills, and knowledge. How these components interact and how quickly each develops weighs heavily on theories of the evolution of our species’ unusual life history and our equally unusual reliance on socially-transmitted behavior. The variation among hunters, both within and among ages, weighs as heavily on models of cooperation and parental investment.

Estimates of hunting skill as a function of age help to constrain and inform theory. A successful theory should predict these functions with an internal logic, or otherwise explain why the data have misled us. And when a model cannot generate observed patterns, it must be either discarded or amended. A detailed model like González-Forero et al. (2017) makes amendments easier because the anomalies have biological meaning. For example, the González-Forero et al. (2017) model predicts that skill increases plateau soon after the onset of reproduction. This may be because learning and production are separate activities in the model. In a generalized linear model, in contrast, a violation has no specific biological interpretation, because the model contains no causal assumptions, only statistical assumptions. While more mechanistic models have advantages over typical generalized linear models, they share many flaws. For example, it is still possible—even likely—that the inferences from data are confounded by factors like cohort effects or the absence of data on components of skill, like ecological knowledge. Combined with an explicit causal model (Pearl et al. 2016), these suggestions guide us to improve future research designs and inferential procedures. So while the work we have accomplished here cannot settle the most important debates, it takes stock of available evidence in a unified framework that will focus and improve future effort.

Our analysis found both a clear central pattern and variation around it. In every site, skill peaks after reproductive maturity. In no society do we find adolescents who regularly display peak individual performance. The average hunter exhibits peak skill around 33 years of age, long after men have reached physical and reproductive maturity. Skill functions are rather flat in the region of the peak, however. Most skill has been achieved by age 18, and declines are typically slow, such that an 80 year old may retain two-thirds of maximum skill. Several sites show peaks that are either much later or much earlier than the average. Among societies such as the Aché of Paraguay (16 ACH, Figure 5), skill increases throughout much of the 30s, peaking only around age 37. This contrasts with sites such as the Matsigenka (9 MTS), where the average peak is at age 24.

Why does skill develop at the observed rates? Are cognitive skills, cultural knowledge, or physical strength to thank? The most plausible answer, based on both data and theory, may be “all of the above.” The average skill function strongly resembles age-related variation in physical strength, both among hunters in subsistence-oriented societies and in modern contexts (Supplemental Figure 12). There are cases where skill arguably peaks after physical strength (Walker et al. 2002), but this does not rule out an important contribution of physical strength to adult skill (Blurton Jones and Marlowe 2002). Adults in their twenties and thirties have also accumulated substantial ecological knowledge (Zent and López-Zent 2004, Demps et al. 2012, Koster et al. 2016). From a theoretical perspective, an optimal life history should develop these components together, with the important caveat that brain growth may need to precede body growth, to enable learning (e.g. González-Forero et al. 2017). Taken together, individual hunters develop physical and cognitive abilities in concert, resulting in high hunting success by their late 20s and early 30s.

The data reveal considerable heterogeneity among hunters. Much of this variation evidently pertains to unmeasured site-level factors, which especially impact the rate at which hunters develop peak skill. On average, for instance, Matsigenka hunters of Peru exhibit their peak more than a decade earlier than Paraguayan Aché hunters, and it is not clear what factors explain such variation. It is common and reasonable for anthropologists to emphasize ecological predictors of cross-cultural variation (e.g. Blurton Jones et al. 1994). But varying rates of skill development may stem as well from mediating social factors that relate only indirectly to ecological differences. Additional theorizing is needed to generate hypotheses about the cross-cultural ontogeny of hunting skill in response to variables such as experience, motivation, opportunities for social learning, and the physical and cognitive demands of hunting in different socio-ecological environments. As opposed to a canalized human life history strategy, this study suggests potential developmental plasticity in traits associated with foraging skill, which manifest not just in contemporary settings but potentially in ancestral settings as well. Therefore, these results imply that singular study sites can rarely be viewed as straightforward analogues for evolutionarily relevant environments (Irons 1998). Regardless of the causes of individual and between-site variation, these differences inform downstream models of risk and sharing economies. Heterogeneity among hunters’ productive abilities alters the effectiveness of food sharing for buffering risk (Boyd 1992). Accordingly, the between-site variation in age-related productivity implies that food sharing has diverse adaptive consequences across sites.

Such questions imply an agenda for future research. Few datasets include longitudinal data on individual hunters, thus hindering analysis of correlations and tradeoffs across the lifespan. Moreover, this study focuses on the hunters’ ages, but hunting returns are expected to vary as a function of traits that covary with age, such as physical strength or ecological knowledge (Gurven et al. 2006). Longitudinal data collection of these and other individual-level variables would permit research on the proximate mechanisms that underlie the skill functions that we have modeled in this study. As a final consideration, our cross cultural data are observational and subject to self-selection biases, implying that potentially hunters enter the dataset only at times when they are expecting to be successful (Heckman 1979). Such biases perhaps explain the divergence of foraging skill across the lifespan, and methodological approaches are needed that explain the representation of hunters in the sample. These are ambitious objectives, but given evidence that subsistence challenges and complexity are a key determinant of life history evolution (Kaplan et al. 2000, DeCasien et al. 2017), a renewed emphasis on foraging skill and production is merited.

## Acknowledgments

We gratefully acknowledge funding from the National Science Foundation (proposal # 1534548). Audiences in London, Lausanne, Nijmegen, Aarhus, UCLA, and the University of Utah contributed useful feedback on draft analyses and interpretations.

## DATA AND CODE ACCESSIBILITY

The data and code used to produce the analyses reported in this paper are available as an R package: github.com/rmcelreath/cchunts.

## SUPPLEMENTAL FILE

**FIGURE 7.**
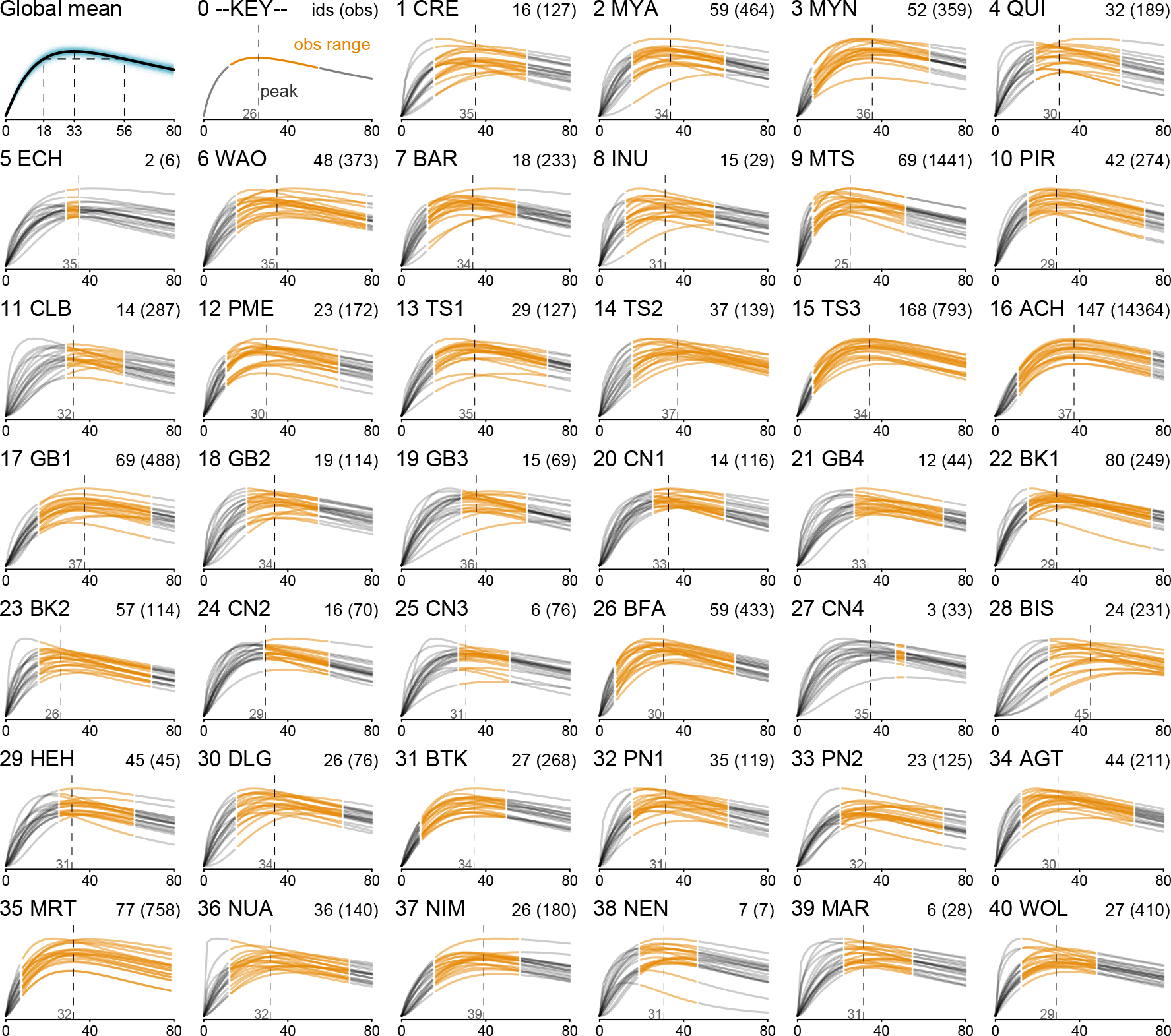
Simulated samples from the posterior distributions of skill functions in each site. This figure is similar to the skill grid in the main text, but it shows simulated hunters, not the posterior means for the observed hunters. This is much better for showing that the model expects more empirical variation than can be seen in the previous skill figure.

### Model definition

Let *y* be an indicator variable for hunting success (produced a non-zero harvest) and *h* any observed non-zero harvest. Let *i* index observed outcomes (harvests). Then:

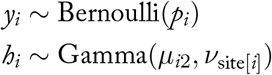

The expressions for *p* and *μ* specify the production functions, indexed by *j* for the outcome type (for successes or harvest size, respectively):

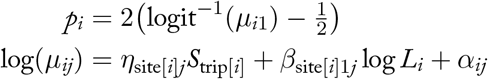

The labor input is *L*_*i*_, the duration of the trip, standardized so that the average trip at each site has *L* = 1.

The skill input *S* into the above is given by the average skill among the individuals contributing labor to a particular observed harvest:

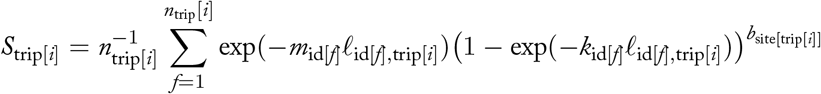

where *n* is the number of productive foragers for trip[*i*] (excluding individuals categorized as assistants, such as porters) and id[*f*] is the forager ID of the *f*-th forager on each trip. This means that for aggregated harvests, in which individual contributions cannot be identified, the model uses average skill. The age *ℓ*_*f*,trip[*i*]_ is the estimated age for forager *f* at the time of trip[*i*]. We describe the age model further down. Note that all ages within the model are standardized by dividing calendar age by the reference age of 80, making *ℓ* = 1 equivalent to 80 years old.

The intercept component of each production function, *α_*ij*_*, is composed from:

- A site-specific intercept *a*_site[*i*]*j*_
- A site-specific and outcome-type specific set of coefficients (elasticities) for the impact of group size, number of assistants, firearms, and dogs. The latter two variables are binary variables indicating whether the hunter had use of a gun (as opposed to other weaponry) or at least one dog.

On the log scale, these combine additively:

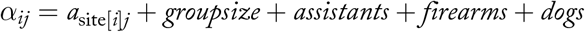

All of these effects are allowed to vary by site as random effects. These assumptions are visible in precise detail in the code to follow.

### Random effects on skill

The life history parameters *k*, *m*, and *b* make use of partial pooling both within and between sites. We use a two-level pooling structure that allows each site to have its own covariance between *k* and *m*. Specifically, let id be the unique ID number of each forager. Then each *k*_ID_ and *m*_ID_ are defined by:

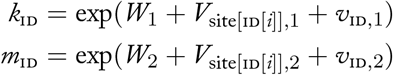

The parameters *W*_1_ and *W*_2_ are overall means, across all sites, and the parameters *V*_*s*,1_ and *V*_*s*,2_ are the offsets of these means for site *s*. This leaves *v*_ID,1_ and *v*_ID,2_ as the offsets for individual ID.

Starting at the lowest level, each pair of parameters v_ID_ = {*v*_ID,1_, *v*_ID,2_} are allocated probability from a bivariate normal:

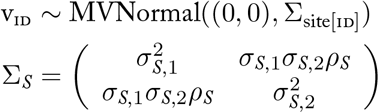

Each site is characterized by 6 parameters: offsets for *k*, *m*, and *b*, as well as standard deviations for *k* and *m* and their correlation *ρ*. These 6 parameters are themselves pooled across sites. This produces the distinction between variance among sites and the variance of the individual hunters, as described in the text.

### Age error model

We accommodate uncertainty in observed ages by defining:

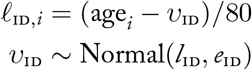

where *l*_ID_ is the observed year of birth and *e*_ID_ is the assigned standard error. In the limit where *e*_ID_ → 0, the age is purportedly known with certainty. Some sites reported ages using uniform intervals. We converted those to Gaussian representations with equivalent variances, so that the imputed ages were unconstrained. In most cases, when a researcher records a uniform age interval, they imply that the true age is closer to the middle of the interval and do not imply that it is impossible for the true age to be outside the interval. To allow this information into the model, we had to use something other than a uniform probability distribution. Gaussian is the most conservative choice, in that case. The irony of the effort put into dealing with age uncertainty is that it has no detectable impact on inference. Fixing all of the ages at their central value produces the same inferences that we reported in the main text.

### Production functions

The skill functions presented in Figure 5 of the main text are inputs into site-specific production functions. These functions have their own elasticities and therefore characteristic shapes. Here we present versions of Figure 5 to illustrate these production functions. There are three different perspectives on the production function. The first component is the probability of success at each age. The second component is the distribution of harvest sizes at each age. These two components multiply to produce the distribution of expected harvests at each age.

To make these components easier to understand, consider all four implied components of the production function for only the Aché sample (Figure 8). The orange functions in the upper-left are the same latent skill functions as in the main text. The red functions in the upper-right are the probabilities of success for each hunter, with the horizontal dashed line showing 50% success rate. The points are the raw data—the proportion of successes at each observed age, aggregated across individuals who were observed at those ages. The lower-left blue functions are the expected harvest sizes, conditional on a non-zero harvest. Again the points are raw data—the average harvest observed at each age. The violet functions in the lower-right are just the product of the red and blue functions, showing the expected harvests at each age.

Each component may be of interest in itself. In some sites, such as the Ache (16 ACH), the success of each hunt contributes more to variation than does the harvest size. The red curves in Figure 8 vary more both across age and across individuals than do the blue curves. As a result, more of the variation in the resulting expected production curves, seen in violet, arises from success rates rather than variation in harvest sizes. The Matsigenka sample (9 MTS) shows the same pattern— more variation in success rates than harvest sizes. This is possibly a result of the prey types available at the respective sites. Regardless of the explanation, decomposing the expected production in this way shows how skill can influence some aspects more than others.

**FIGURE 8.**
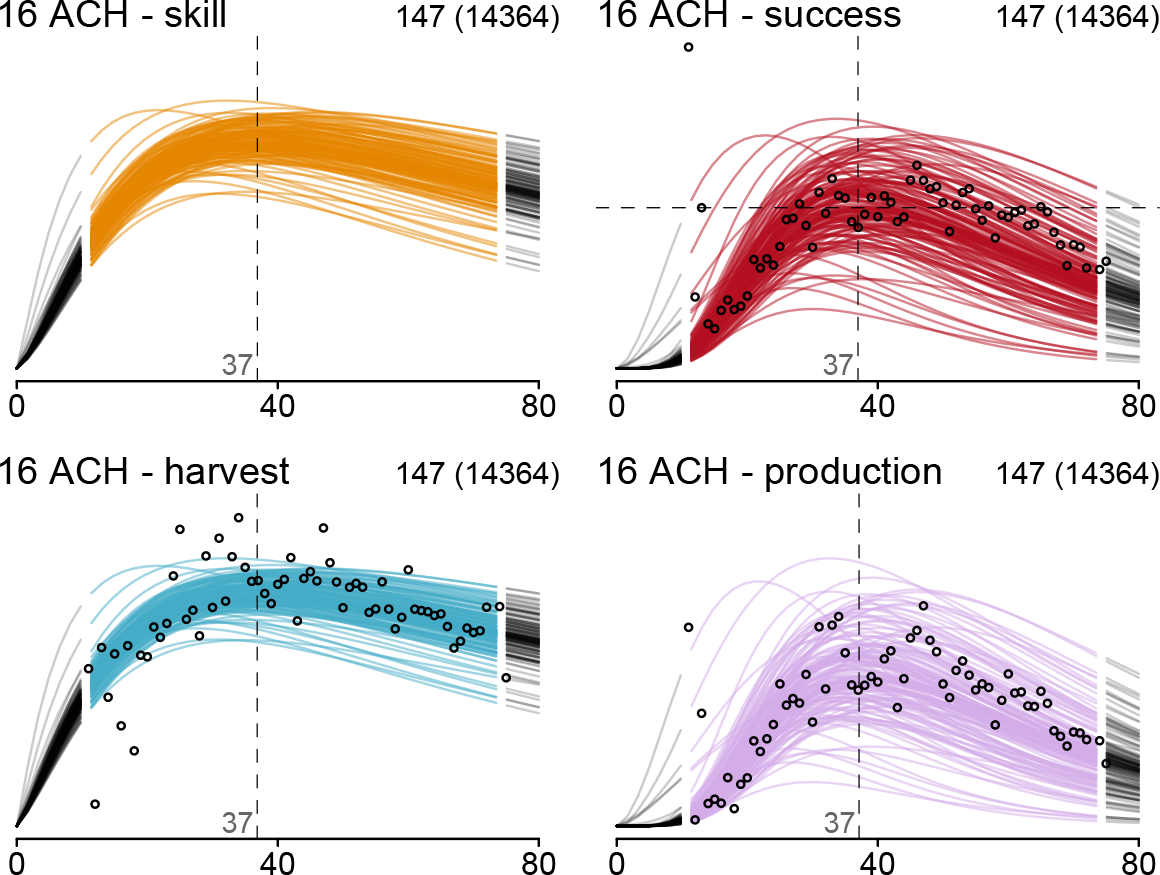
Components of the forager production functions for the Aché sample. See text for description.

**FIGURE 9.**
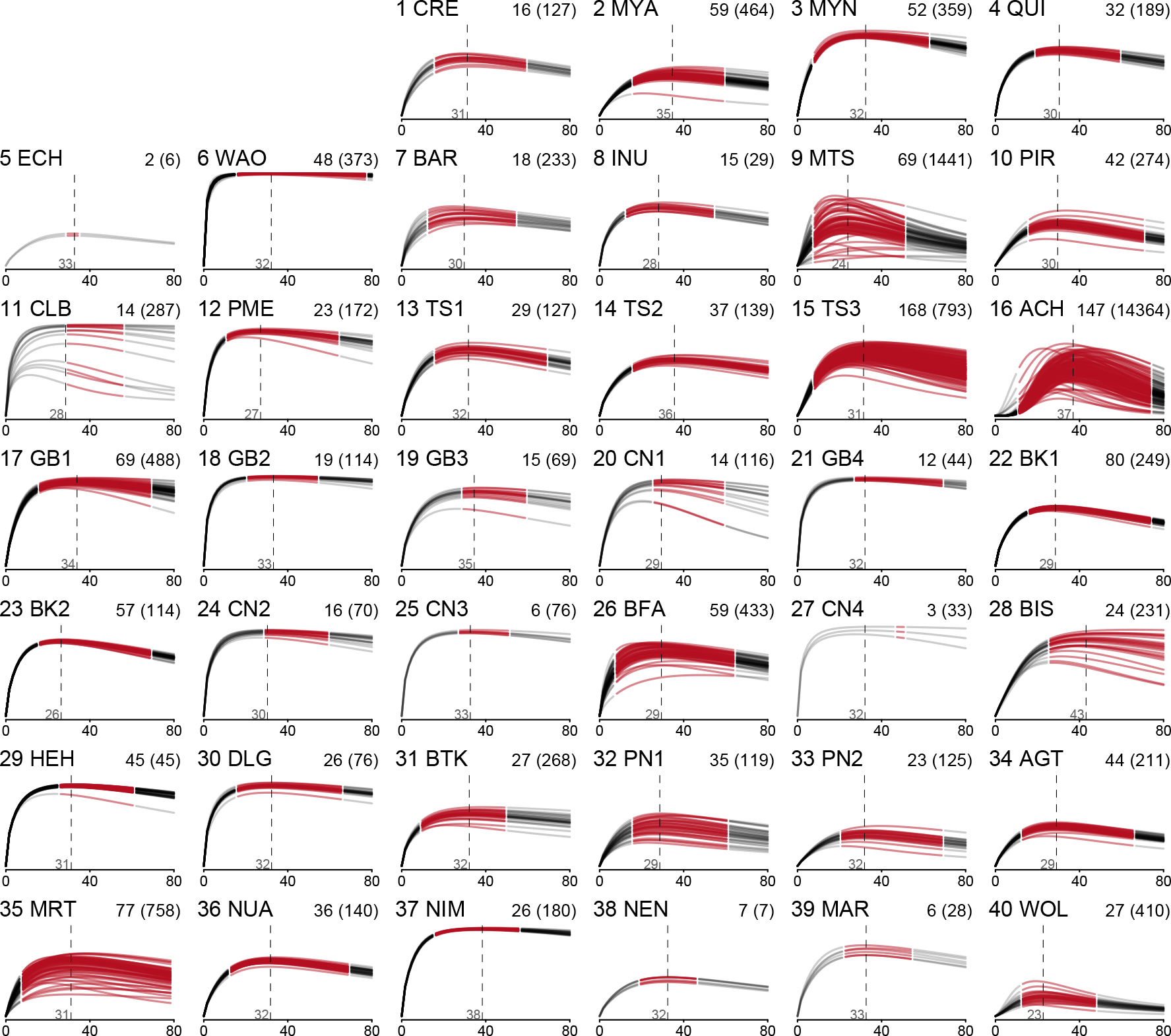
Posterior mean probabilities of hunting success across age. The axis ranges from 0 to 1, and is the probability of hunting success (a non-zero harvest). Several sites, such as GB4 (21) and DLG (30), show essentially no variation in hunting success, since virtually all documented trips result in a non-zero harvest. Other sites, such as MRT (35) and WOL (40), show substantial failure rates and variation arising from it. nb: Variation in methods for documenting unsuccessful hunts imposes limitations on comparisons across sites.

**FIGURE 10.**
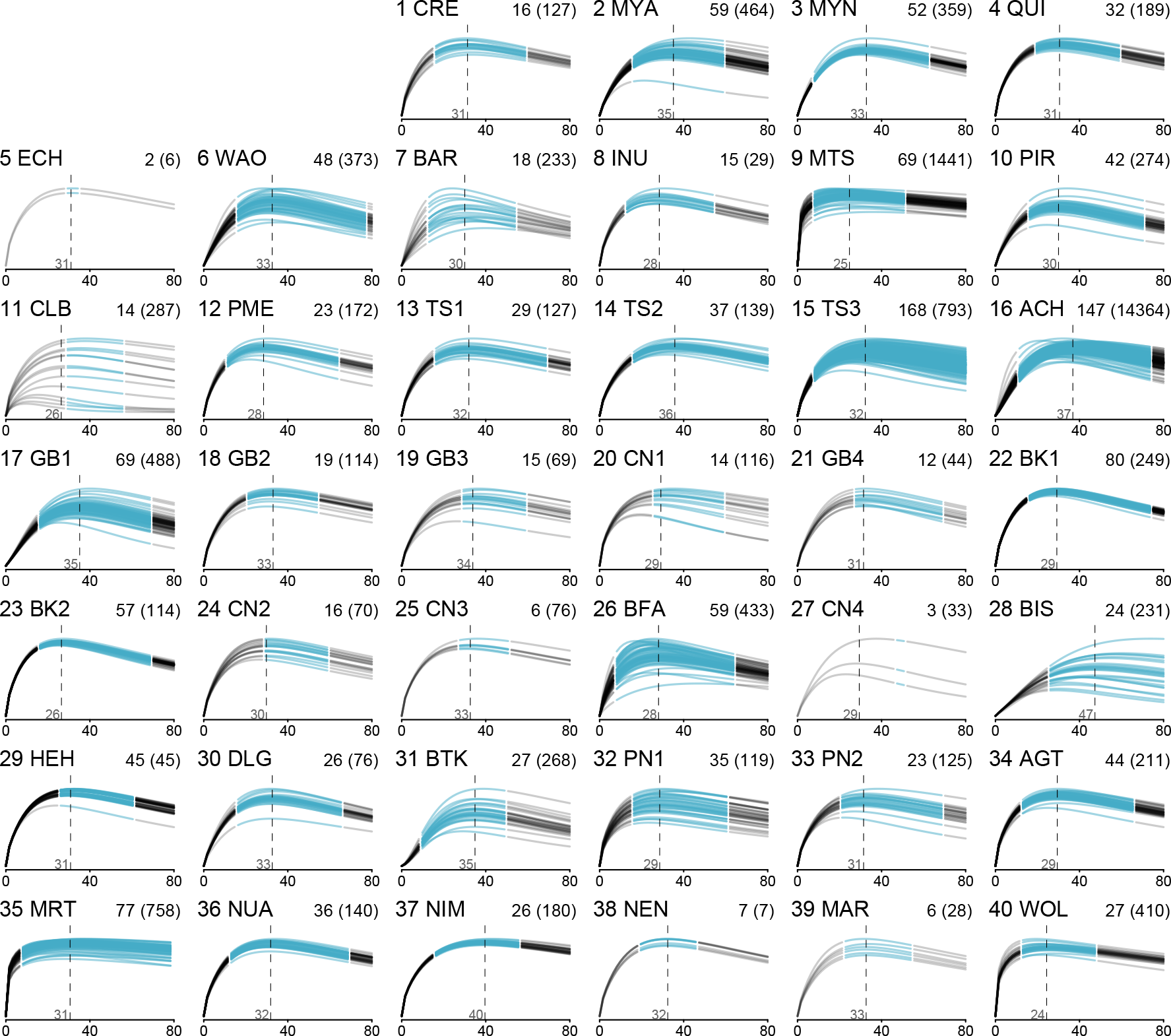
Posterior mean non-zero harvest size across age. The vertical axis is proportion of maximum harvest at each site. So while the units are uninformative, variation remains interesting.

**FIGURE 11.**
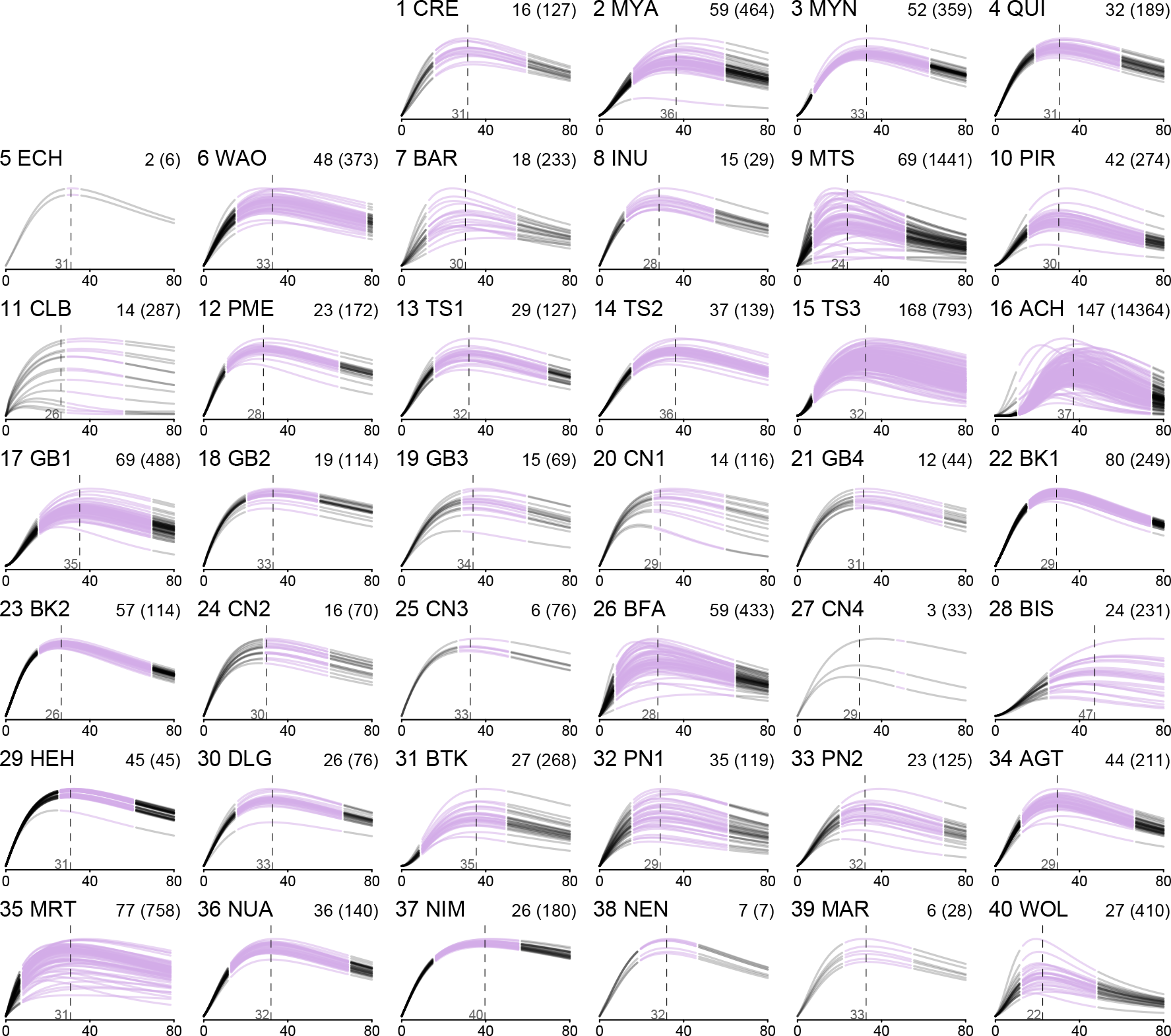
Expected production across age. These functions are just the product of the success function and the expected harvest function. In considering relative expected energy contributions of individuals at different ages, these curves are perhaps the most relevant representations of the data.

**FIGURE 12.**
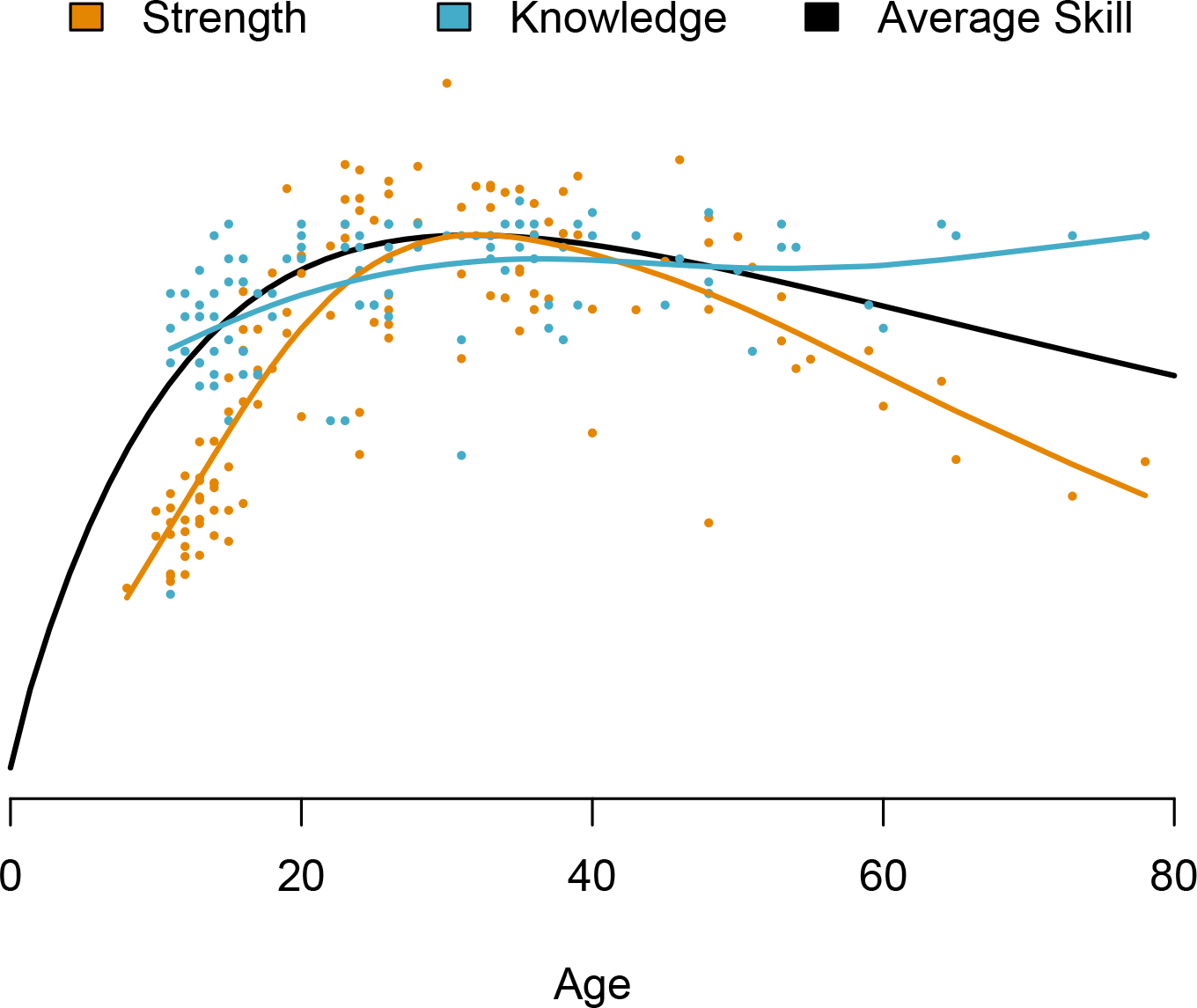
Strength and ecological knowledge as a function of age. Points depict the strength and knowledge of males at the lead author’s field site in Nicaragua. Strength is measured using methods described by Gurven et al. (2006). Knowledge reflects performance on questions about fish behavior, as described by Koster et al. (2016). Fitted lines reflect smoothing splines, and they are superimposed atop the average skill function from Figure 5. All fitted lines are standardized to have the same maximum value, but knowledge has an expected minimum of 50% of the maximum, reflecting the guessing probability of the binary questions. Note that there is minimal residual correlation (*ρ* = 0.09) between individual men’s knowledge and strength after partialling linear and quadratic effects of age from the respective response variables.

### Detailed model code

The code for the model is available in the accompanying R package. In this section, we explain the model block of the code, focusing on how the marginalization over missing values is accomplished.

The first portion of the model block defines local variables, used in calculations, and priors. The only unusual code here is the Jacobian adjustment applied to `lifehistmeans[4]` and `lifehistmeans[5]`. This adjustment allows us to apply the prior on the natural, instead of logarithmic, scale.

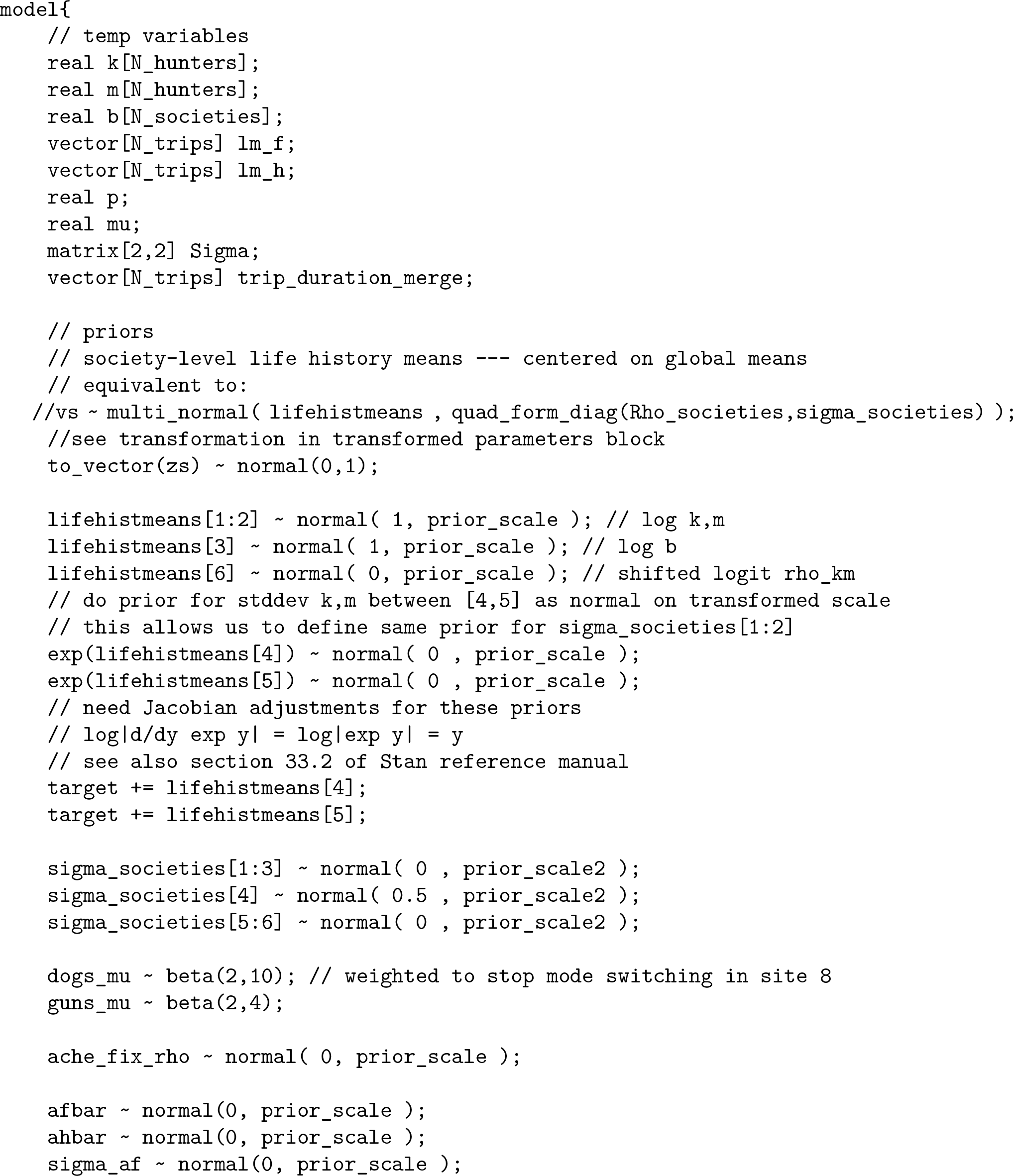

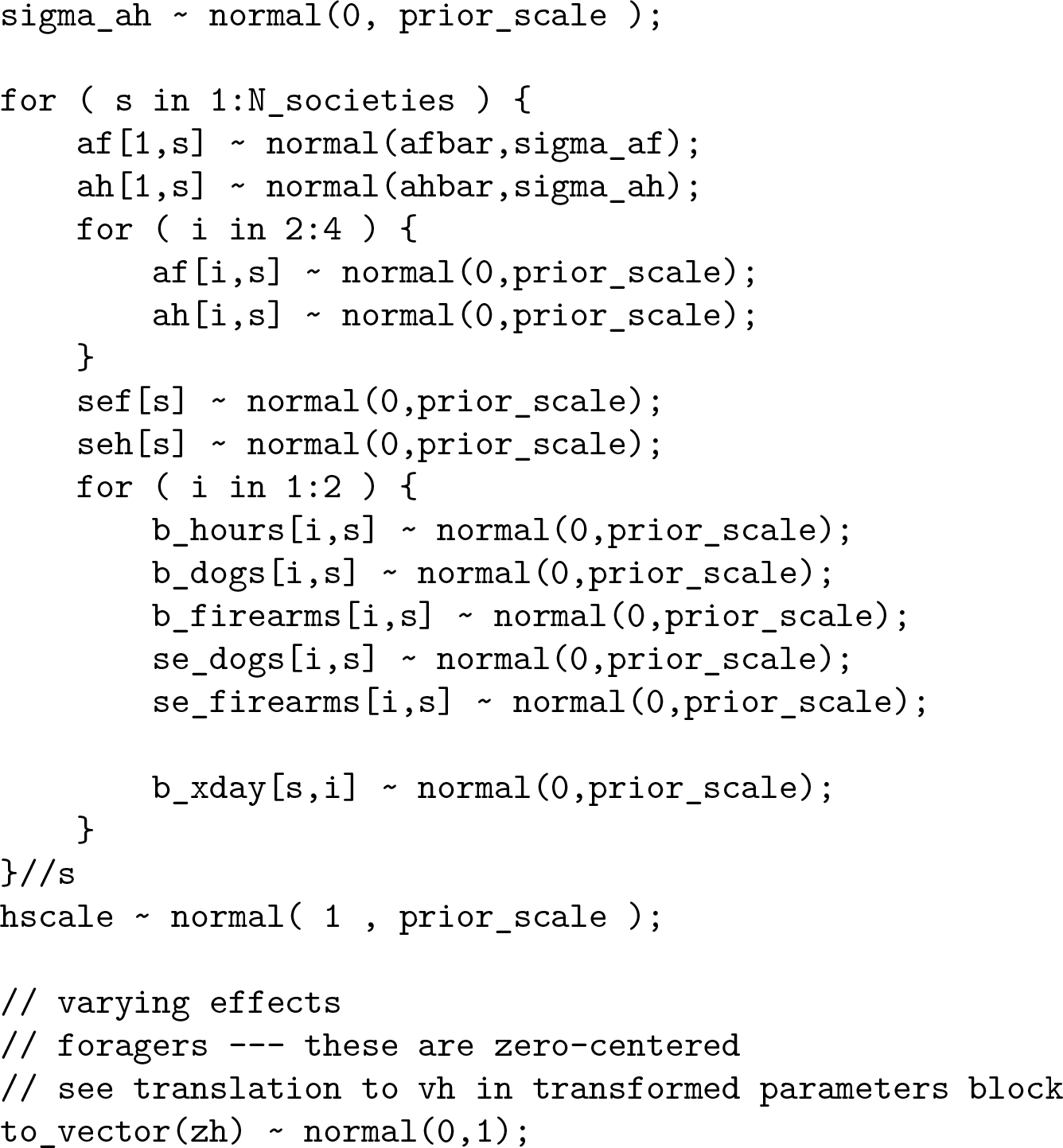

The next chunk of code handles imputation of missing ages and trip durations. For each missing age, there is a corresponding standard error of the age. This comprises a Gaussian prior for the error of each missing age. Combined with the prior for each missing age, this provides a way to average over the uncertainty. For each missing trip duration, similarly a parameter is used. Then a vector that merges observed and missing values is generated. The prior formed from each site’s (standardized) trip durations constrains the imputed values.

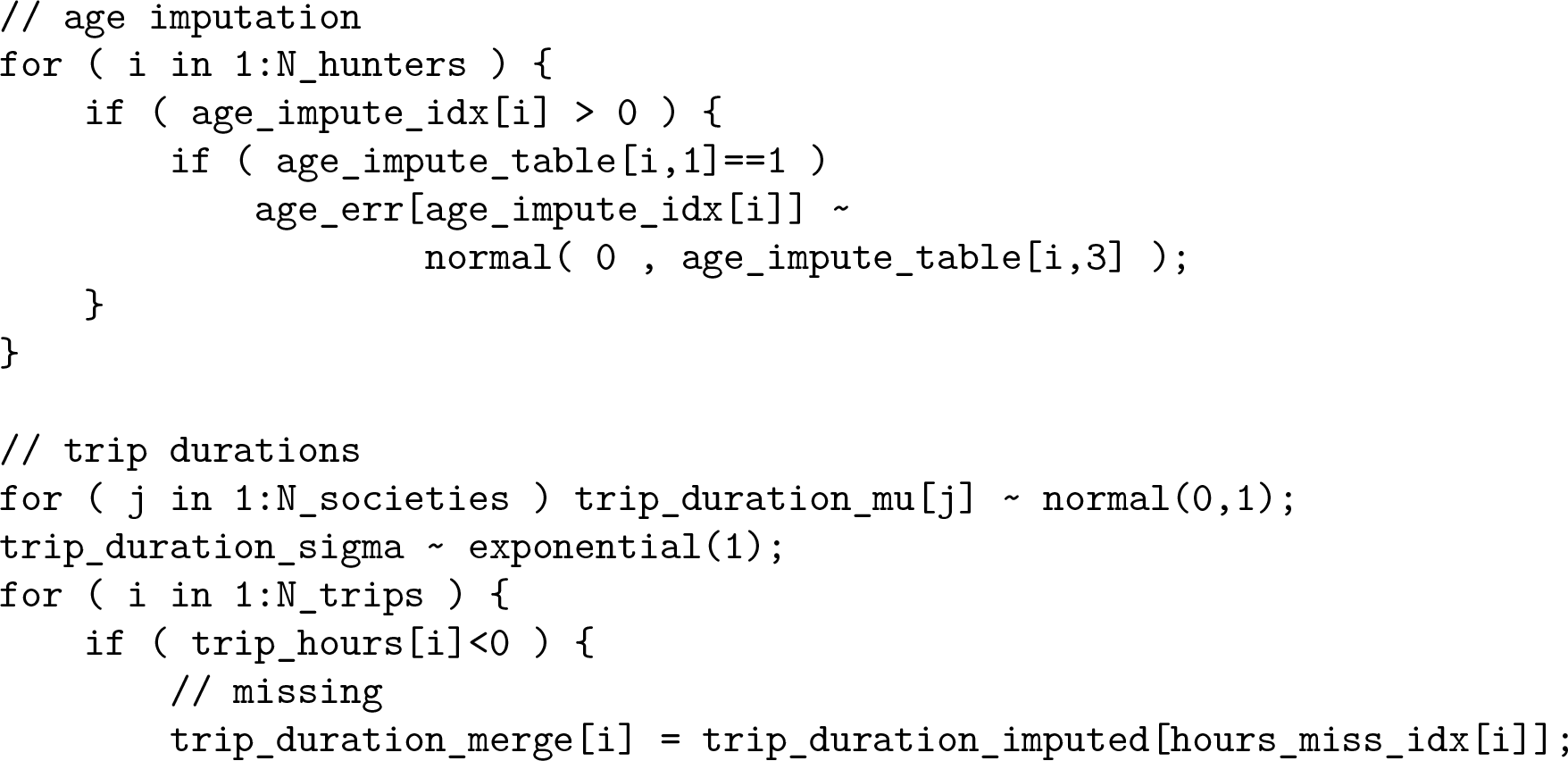

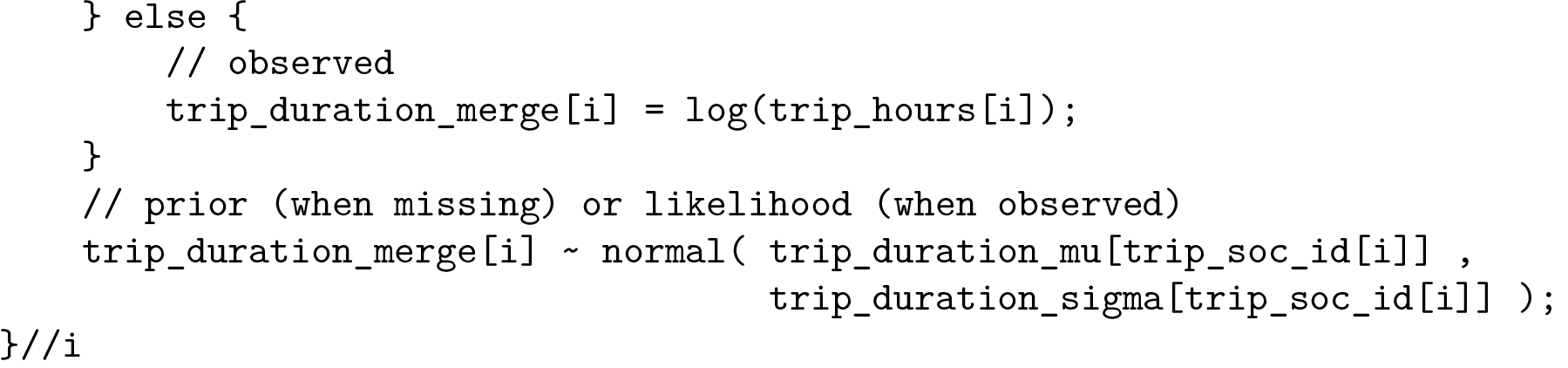

The next short section computes hunter-specific and society-specific skill parameters. These are then reused in the likelihood calculations to follow.

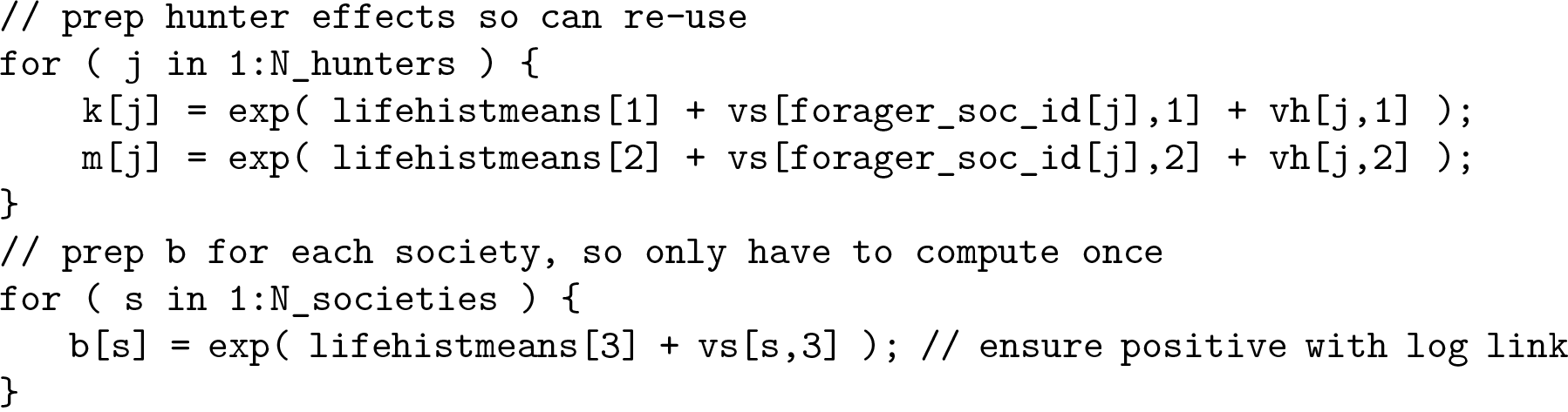

The main loop of the model block comes next. This loop passes over trips, and then harvests within 750 trips. The first chunk of code just prepares local variables. The `xdogsvec` and `xgunsvec` arrays exist to help us construct marginal log-probabilities when both dogs and firearms are unobserved (missing). The relevant code appears later down.

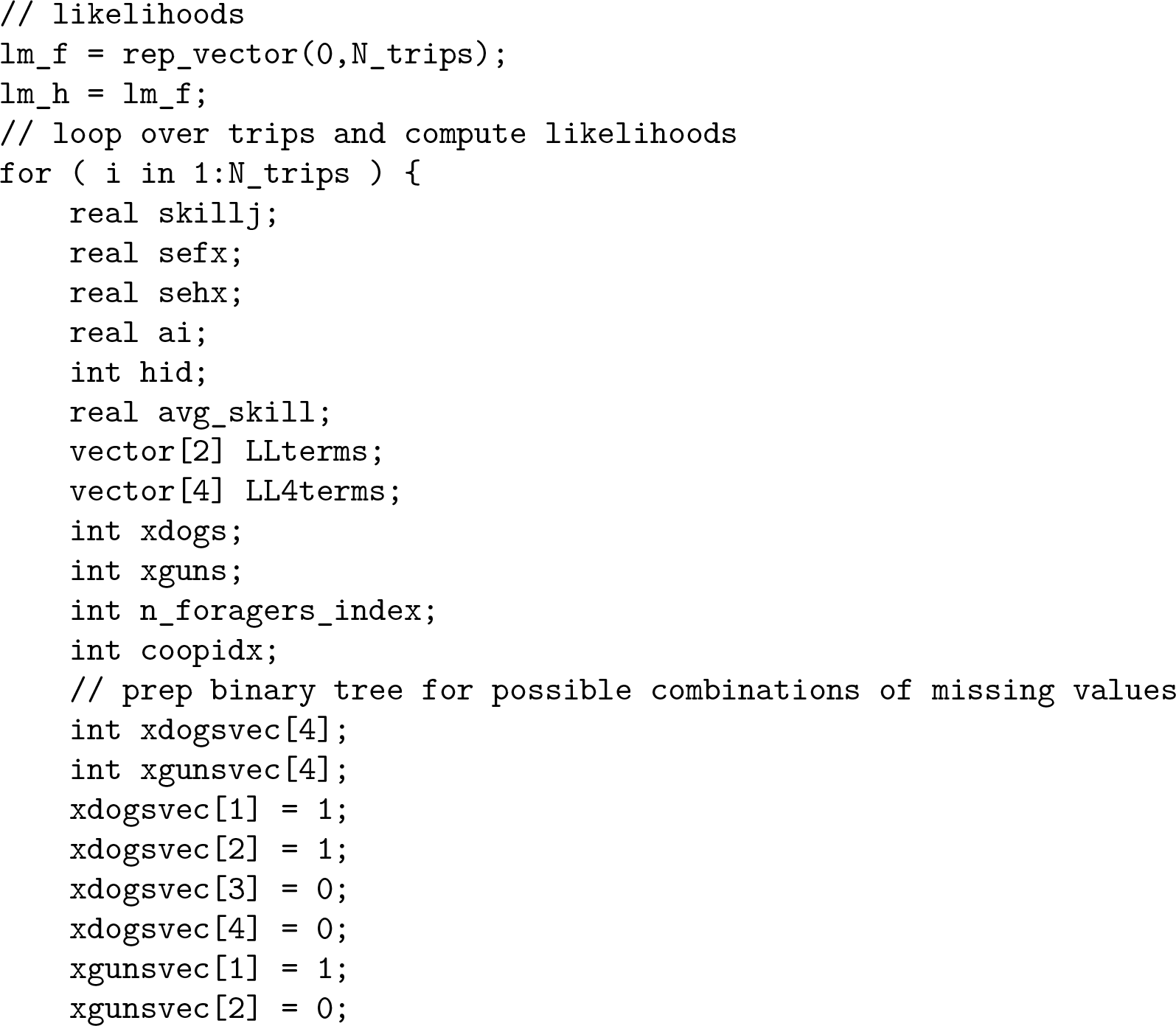

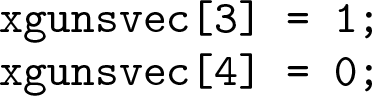

Next, when a trip has a pooled harvest, average skill for the entire group of hunters must be calculated. This is because we assume that production depends upon average skill in this case, where we cannot identify individual contributions. The `coopidx` variable tells us later which intercept parameter is needed, as the intercept in production differs depending upon pooled or individual harvests.

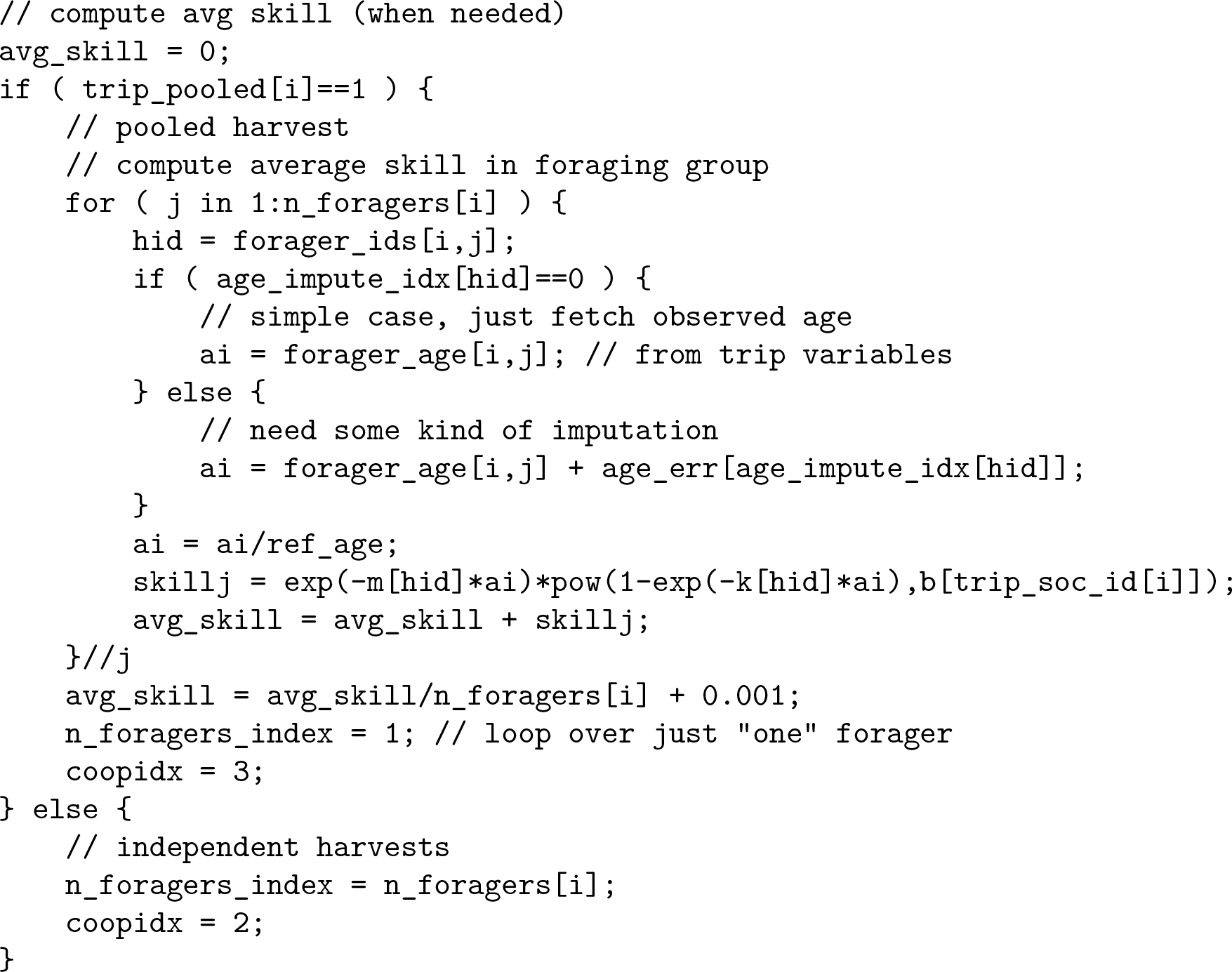

The big loop over individual foragers comes next. The loop begins by calculating individual forager skill, but only when harvest is not pooled. This code is structural the same as that used above to compute average skill, but it omits the averaging.

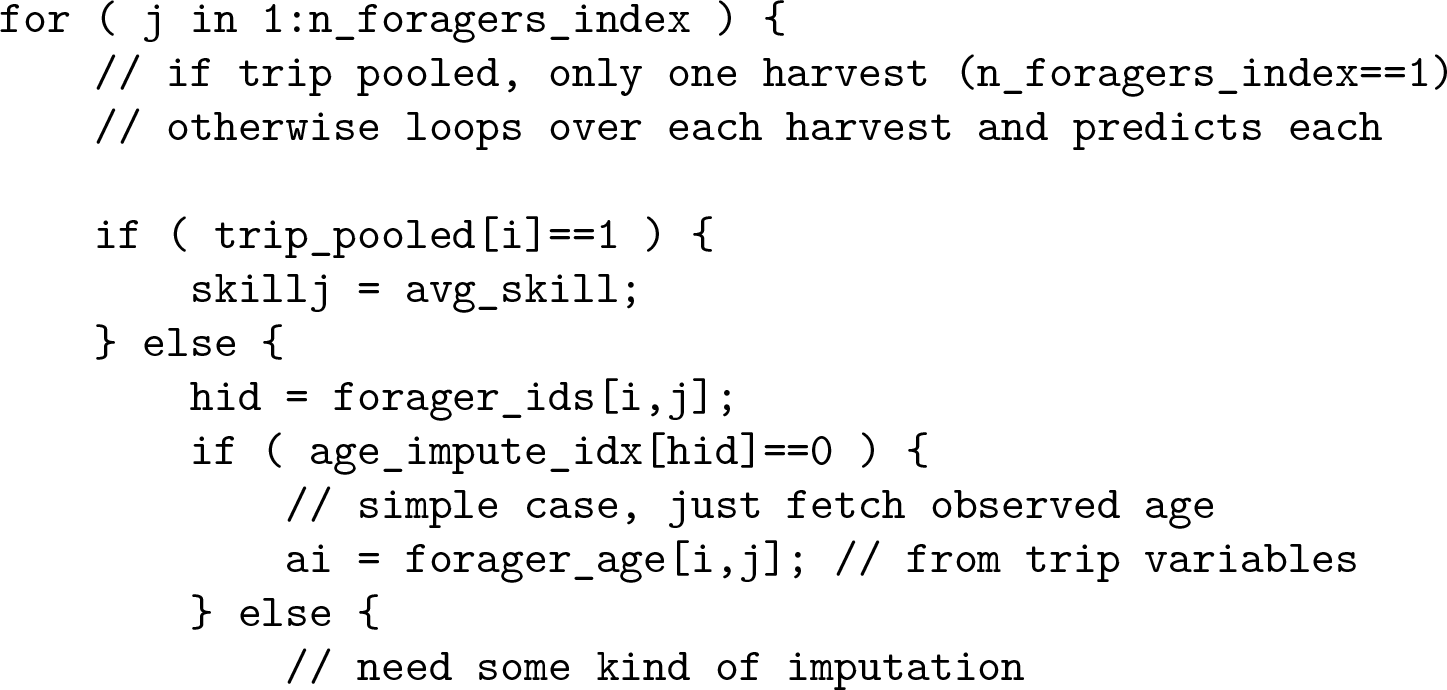

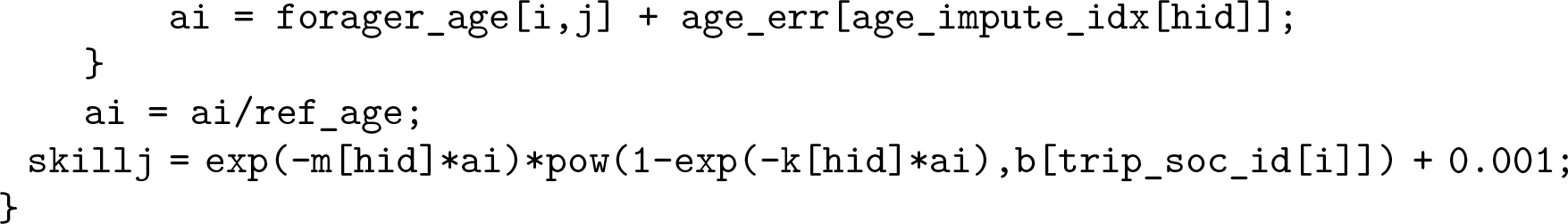

Next we build “stem” expressions for each harvest log-probability. These stems contain all terms except those for dogs and firearms. Dogs and firearms must be added conditional on missingness. In that case, these stems are reused for each missingness state.

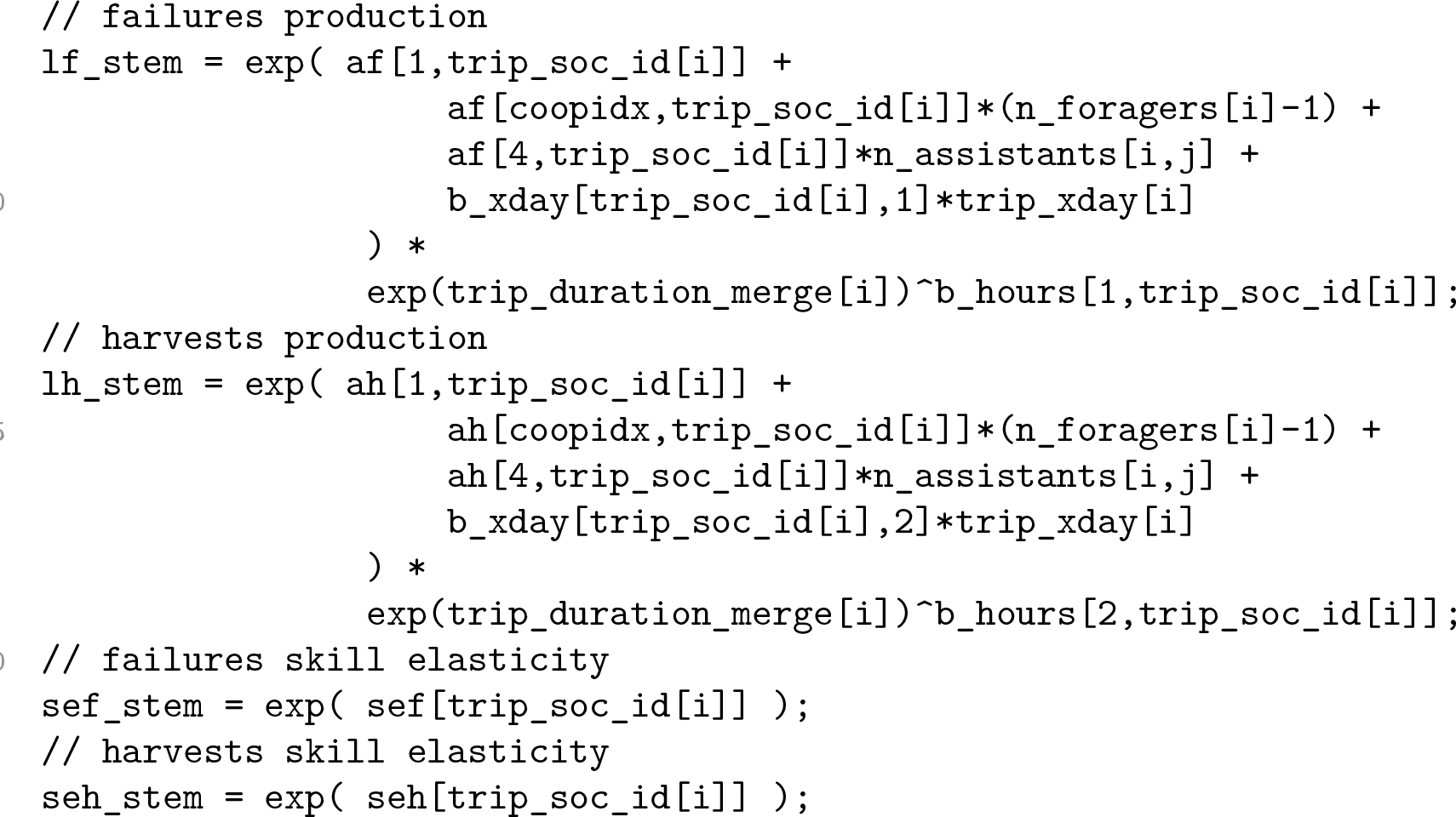

Now we can do target updates. Different expressions need to be built, depending upon whether dogs, firearms, or both are missing. The simplest case is when both are observed. In this case, we just add the observed values to the stems, compute probability of failure, average harvest, and update. Note that *−*1 as the missingness indicator is chosen during data initialization. Note that the code here considers the probability of a zero harvest, instead of the probability of a non-zero harvest. This is equivalent to the analytical model definition given earlier, even though the expression looks different.

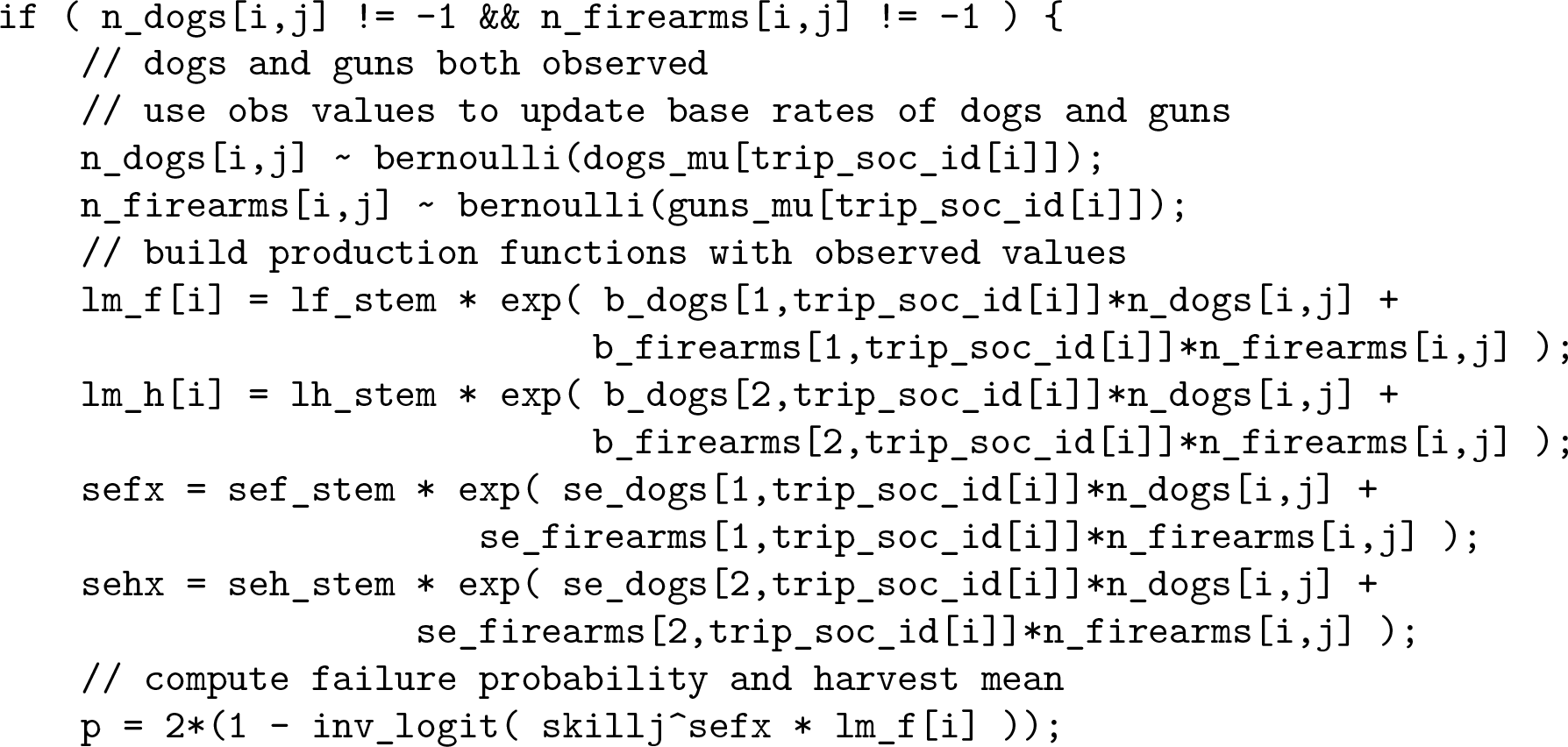

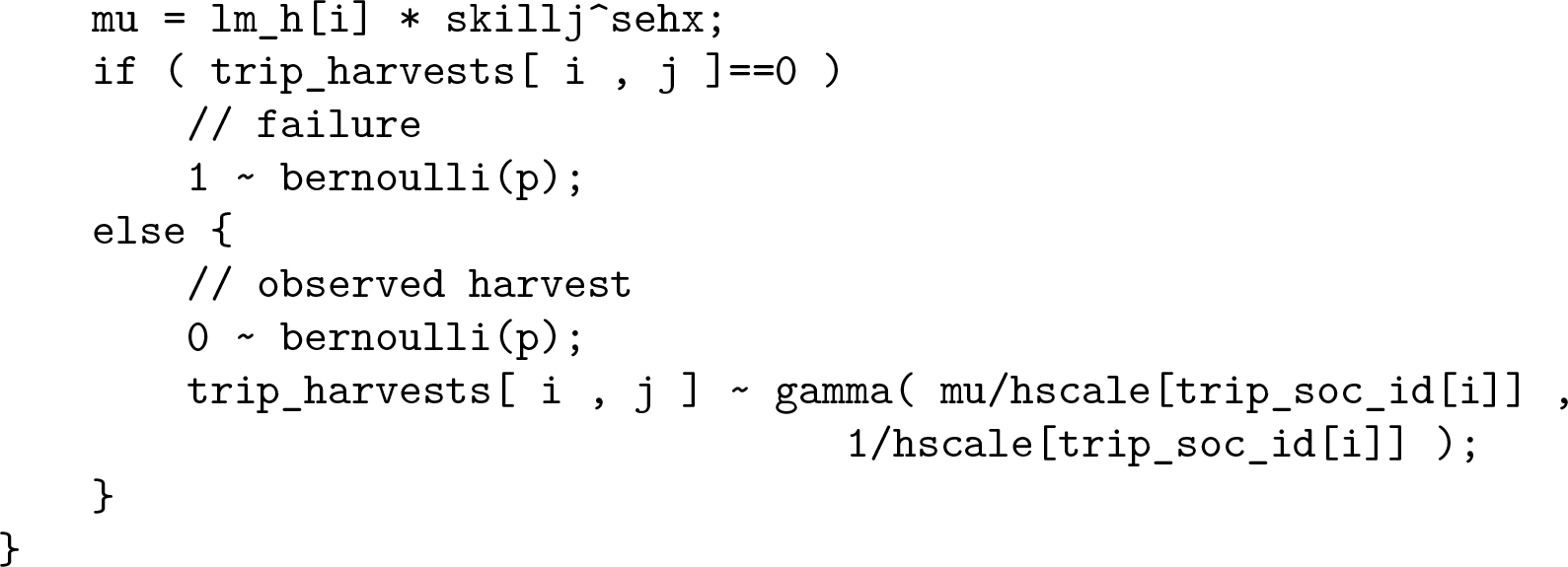

The next two cases are when either dogs or firearms are missing. In these cases, we need to marginalize over missingness states. This generates two log-probability terms in a mixture.

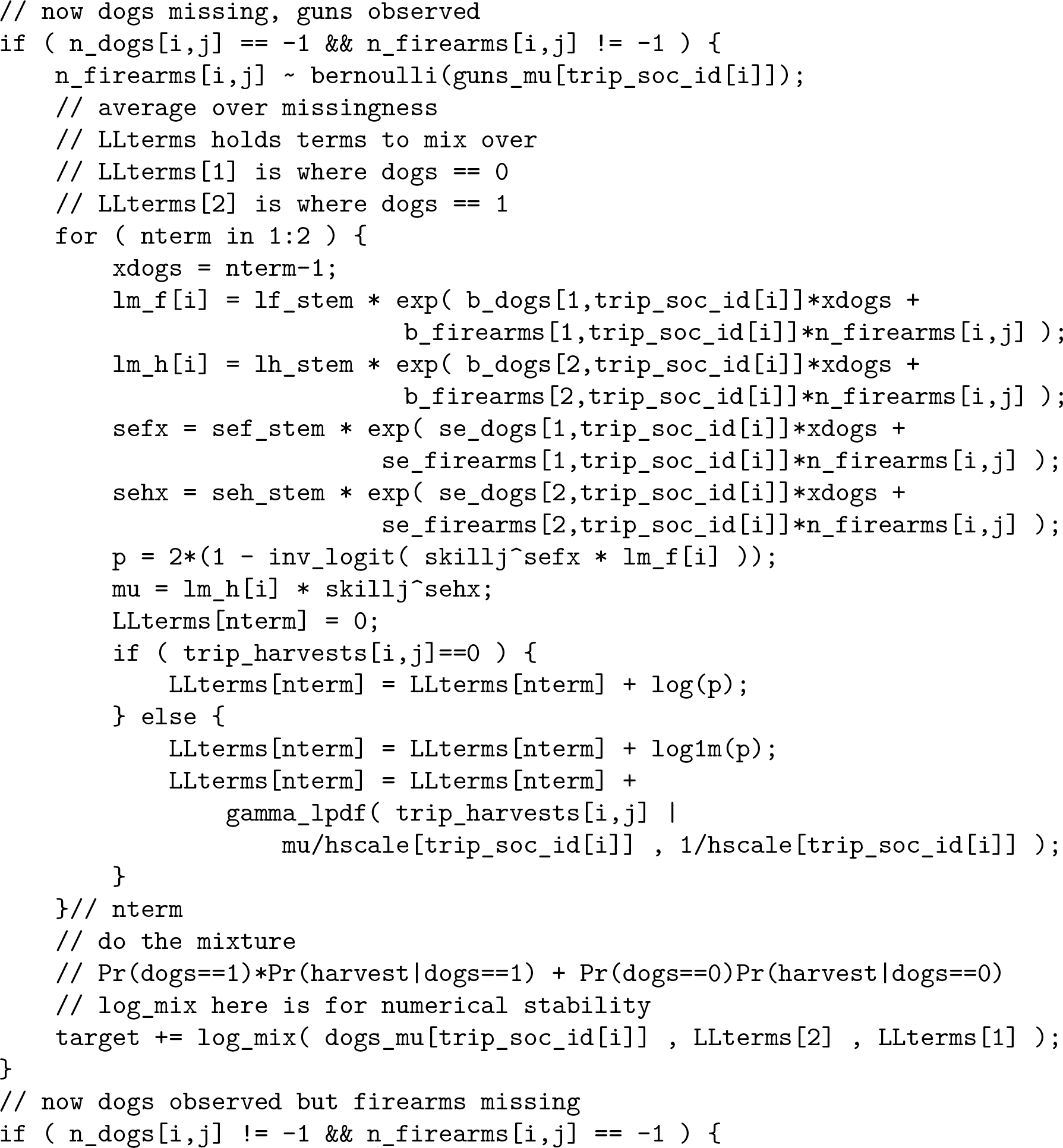

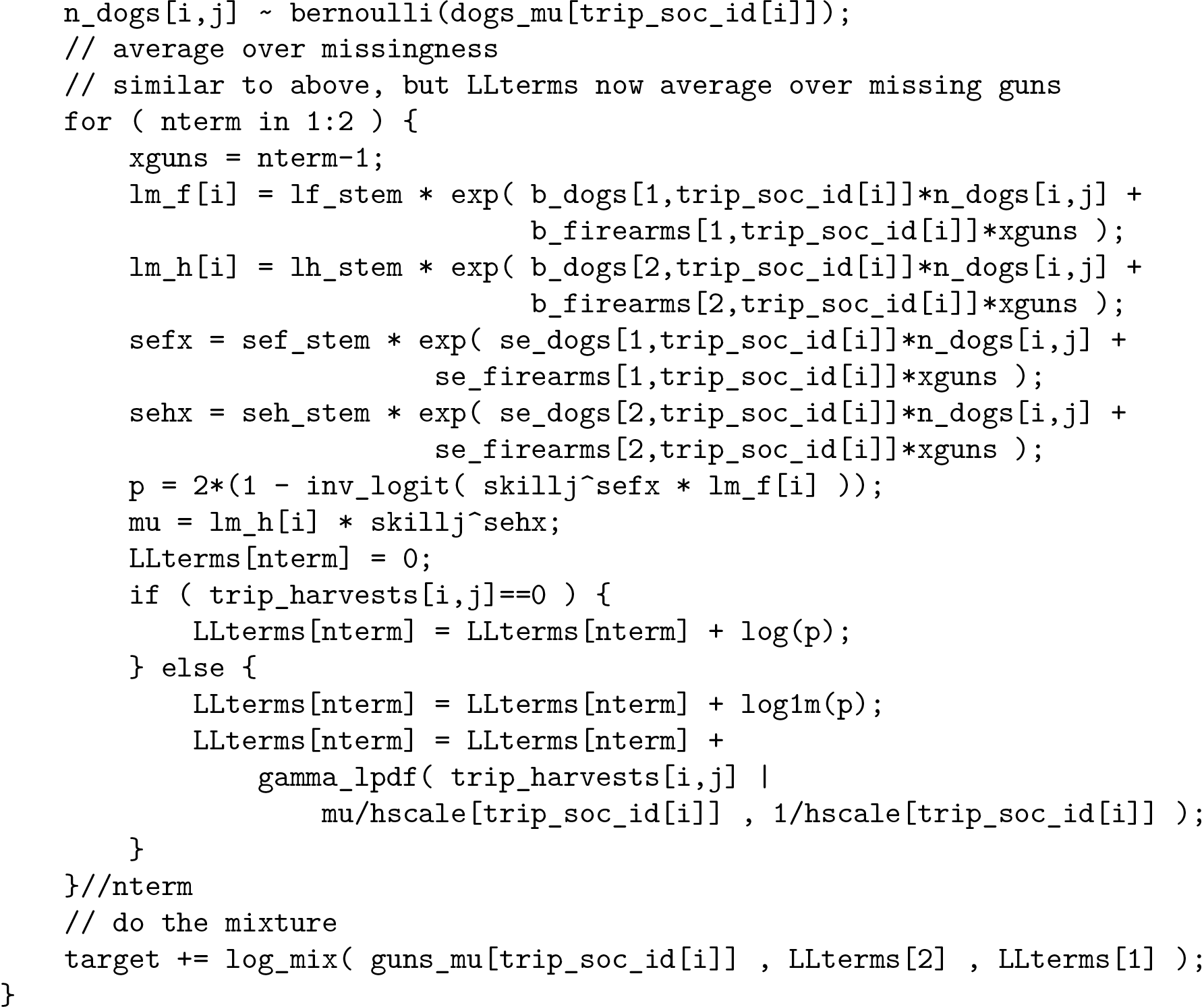

Finally, both dogs and firearms could be missing. In this case, we need a mixture over four possible states.

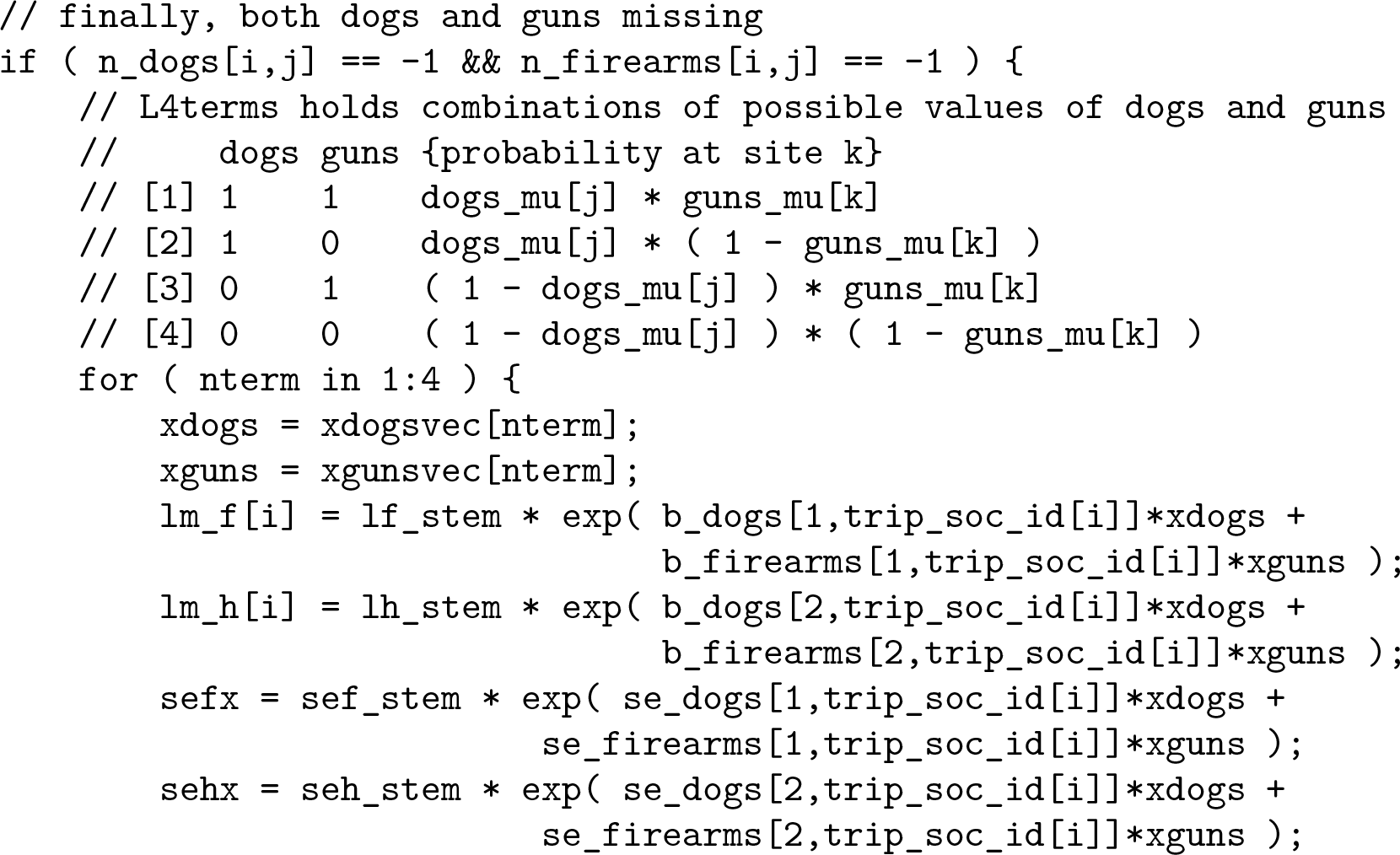

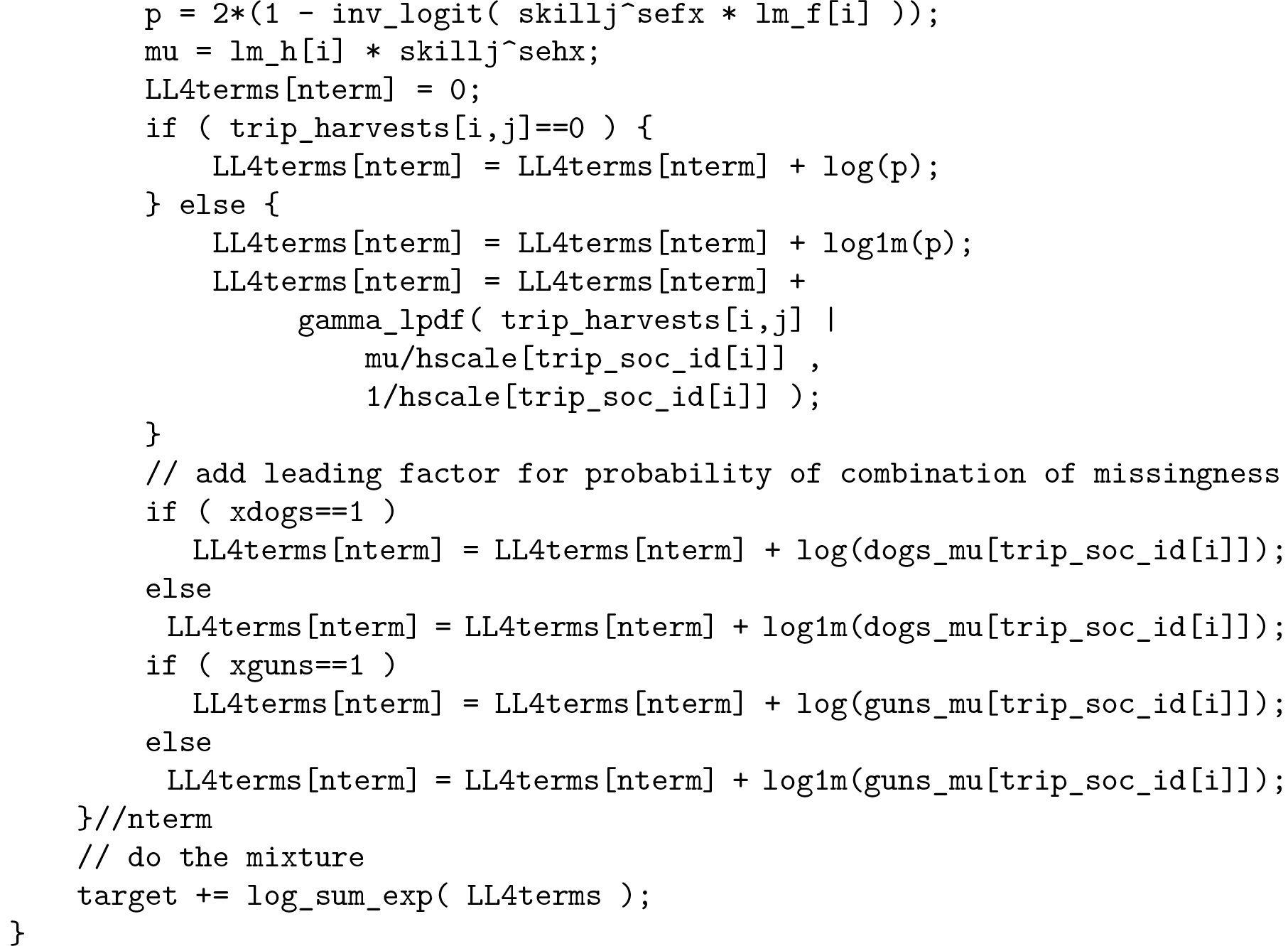

In the end, the model block just loops over foragers j and trips i until all trips have been processed.

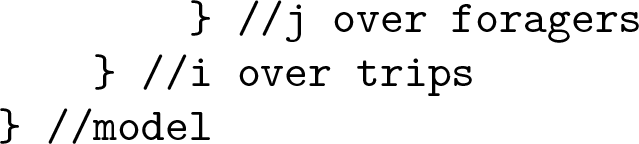

### Model robustness

We went through a lot of effort to handle age uncertainty, missing values in trip durations, and missing values in technology (dogs and firearms). Models that ignore these issues produce very similar inferences for skill functions. On the one hand, this is disappointing, because it really was not trivial to do the right thing, and it did not seem to matter much. On the other hand, it is important to do the right thing, even if it turns out not to matter.

### Marginal posterior distributions

Many of the parameters in the production functions are interesting in themselves. For example, the marginal effects of group size and technology inform debates about human economies. In the figures that follow, we present marginal posterior distributions for all of these parameters, labeled informatively.

**FIGURE 13.**
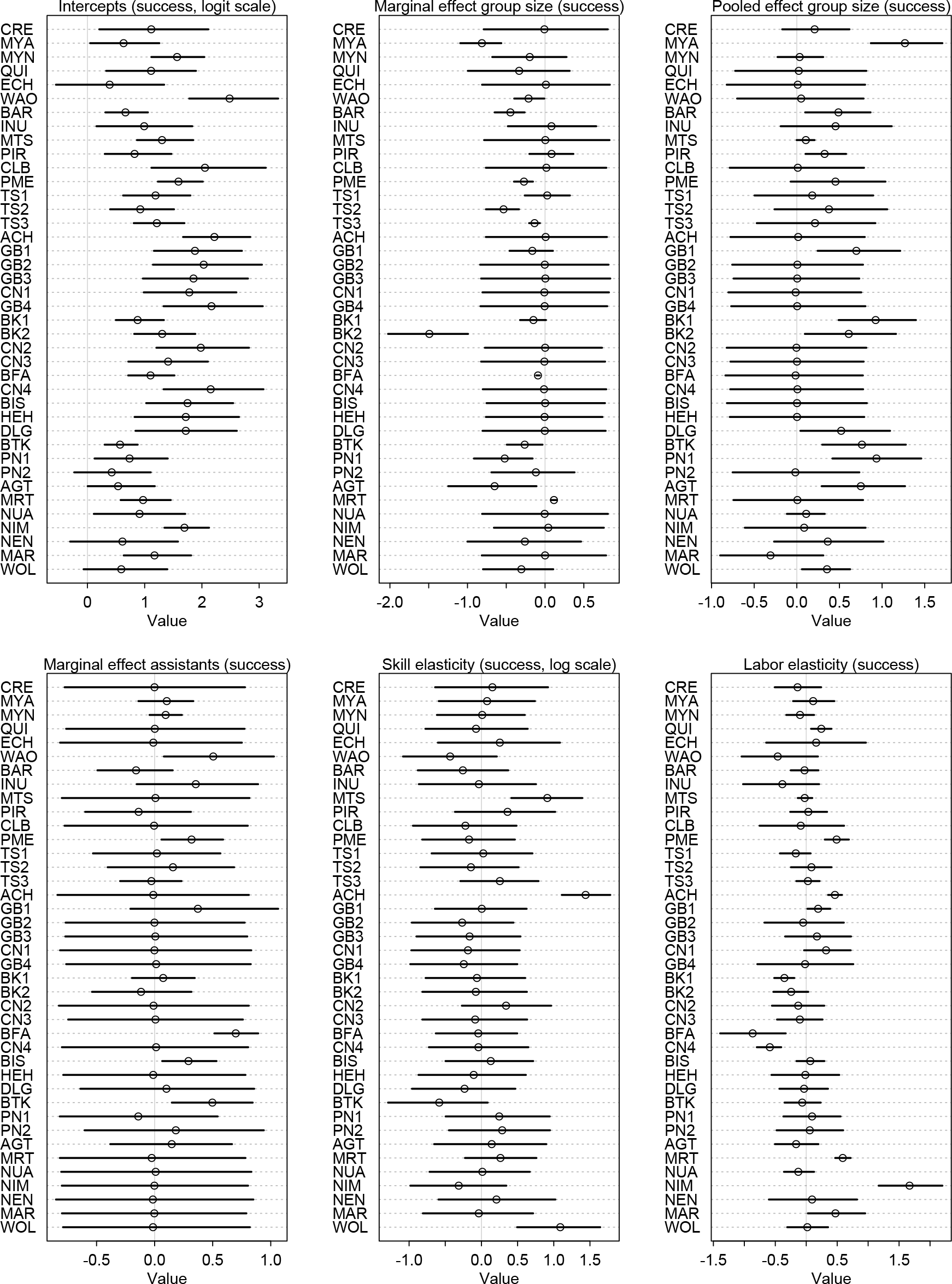
Marginal posterior distributions for production components (success). In the code, these parameters are named `af[1], af[2], af[3], af[4], sef`, and `bhours[1]`, respectively. Note that marginal distributions centered on zero with standard deviation 0.5 correspond to the prior. In those cases, the society contained no information to inform the parameter.

**FIGURE 14.**
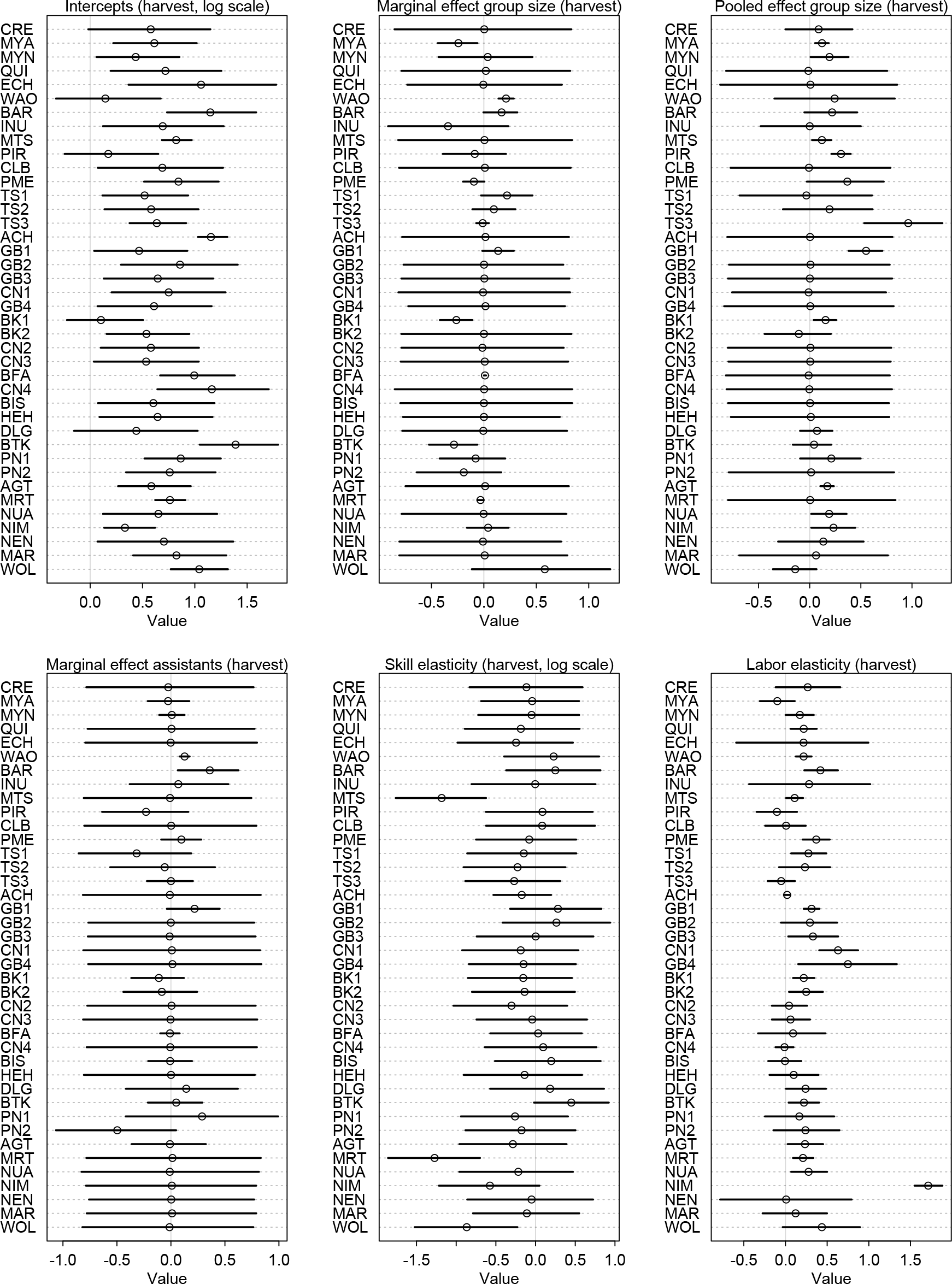
Marginal posterior distributions for production components (harvest). In the code, these parameters are named `ah[1], ah[2], ah[3], ah[4], seh`, and `bhours[2]`, respectively. Note that marginal distributions centered on zero with standard deviation 0.5 correspond to the prior. In those cases, the society contained no information to inform the parameter.

**FIGURE 15.**
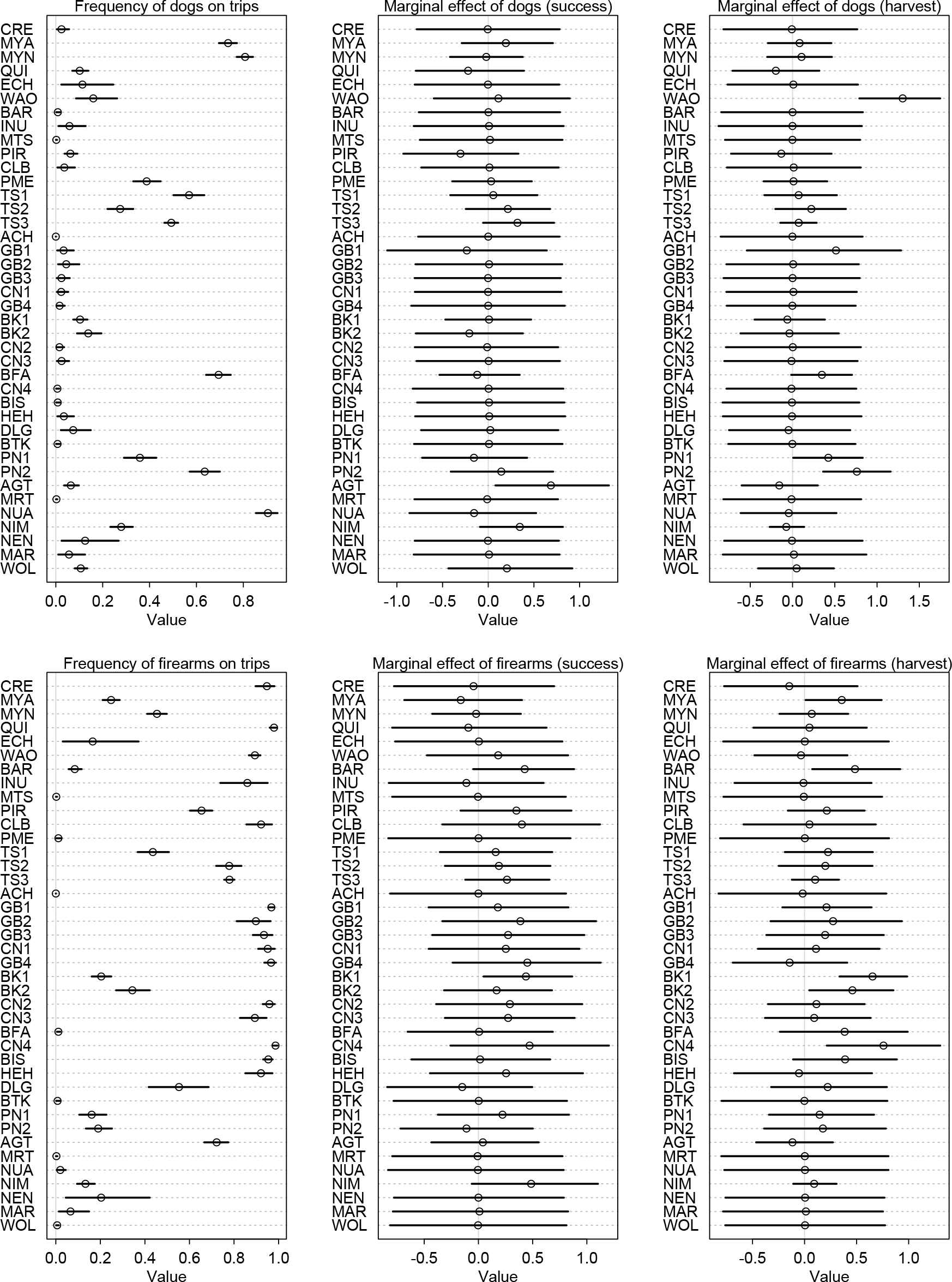
Marginal posterior distributions for dogs (top row) and firearms (bottom row). In the code, the separameters are named `dogs_mu,bdogs[1], bdogs[2], firearms_mu, bfirearms[1]`, and `bfirearms[2]`, respectively. Marginal distributions centered on zero with standard deviation 0.5 correspond to the prior. In those cases, the society contained no information to inform the parameter. Dogs are used at two sites, MTS and HEH, in which their use on trips was not documented. These missing data were averaged into the intercept and set to zero in this figure.

**FIGURE 16.**
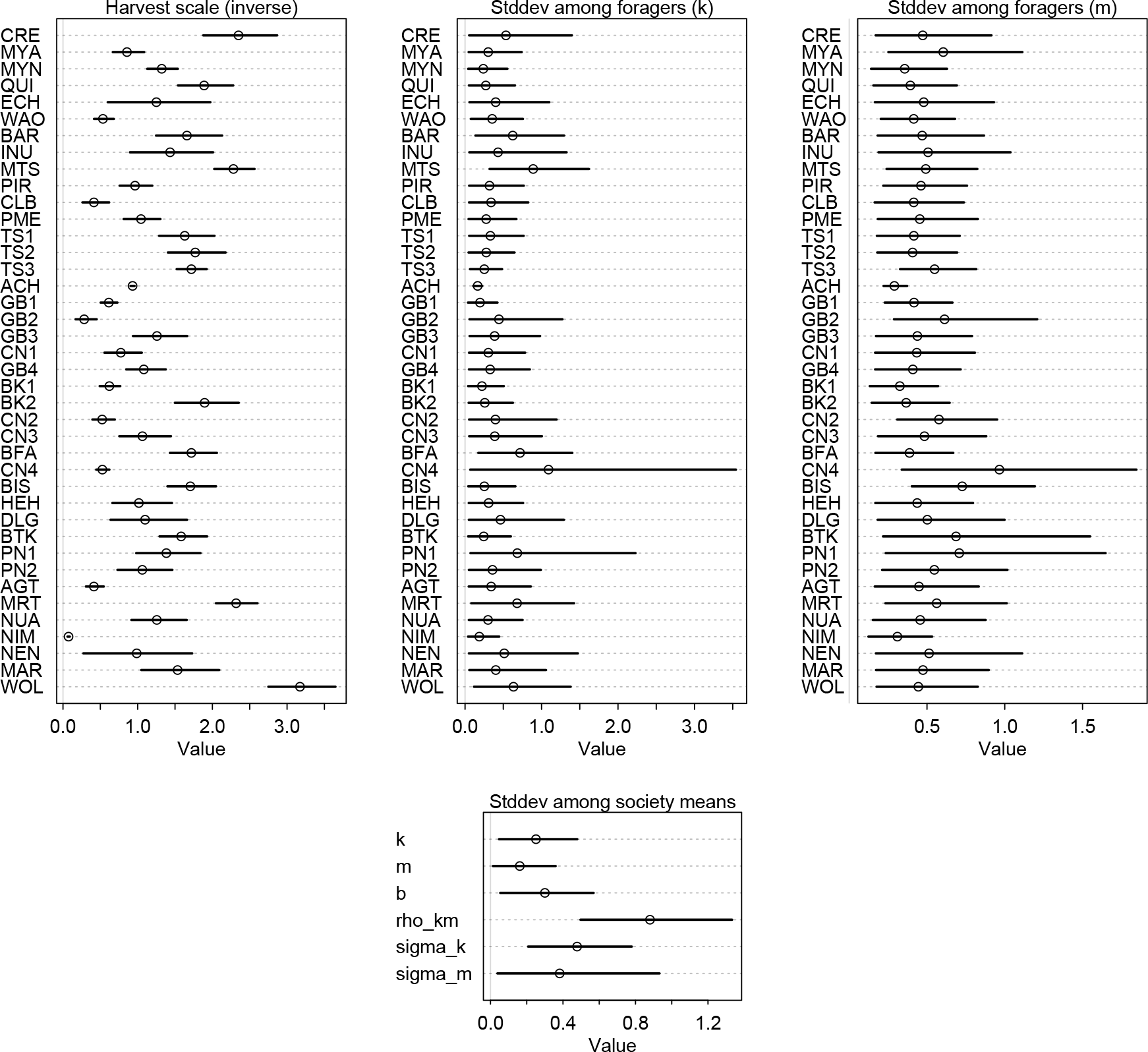
Marginal posterior distributions for dispersion and variance components. In the code, these parameters are named `hscale` (top-left), `sigmas_hunters[1]` (top-middle), `sigmas_hunters[2]` (top-right), and `sigma_societies` (bottom-middle). In the bottom-middle, *k* indicates the standard deviation among sites in mean skill growth, *m* the standard deviation among sites in mean skill decay, *b* the standard deviation in *b* across sites, `rho_km` the standard deviation (on the latent scale) of the correlations between *k* and *m* across sites, and then `sigma_k` and `sigma_m` are standard deviations across sites of standard deviations among foragers in each site.

## Footnotes

1 https://osf.io/2kzb6/?view_only=682fcab2dd614dbdb015612b83044f49

